# Evidence of tornadic phenomena in cerebral aneurysms

**DOI:** 10.64898/2026.08.07.743435

**Authors:** Valentina Mazzi, Diego Gallo, Thangam Natarajan, Jonas Schollenberger, Karol Calò, David Saloner, David A. Steinman, Umberto Morbiducci

## Abstract

Cerebral aneurysms are abnormal outpouchings of arteries within the brain and occur in ∼1 in 30 adults. Their initiation, growth, and rupture have been linked to focal blood flow abnormalities—often termed “disturbed” or “hostile” hemodynamics—but commonly-used hemodynamic metrics yield conflicting associations with pathology and lack a unifying mechanistic interpretation. Building on a theoretically-grounded link between wall shear stress and near-wall vorticity, we hypothesized that a topology-based description of near-wall flow can operationalize the concept of hostile hemodynamics in a reproducible way. Inspired by atmospheric tornadic phenomena, we sought a principled taxonomy of coherent near-wall fluid structures with potential mechanobiological and clinical implications. Using high-fidelity computational fluid dynamics simulations in anatomically realistic geometries, we identified coherent near-wall fluid structures whose organization mirrors well-studied atmospheric phenomena: tornado-like columnar rotating cores; downburst-like nonrotating wall-impinging jets with tangential outflow, roll-cloud-like tangential vortices; and mixed configurations. These tornadic events on the aneurysm luminal surface were identified from wall shear stress topology, consistent with its theoretical connection to near-wall vorticity kinematics. The presence of tornadic phenomena—and their imprints on the aneurysm wall—was independently observed *in vivo* using 4D flow magnetic resonance imaging. By translating concepts from atmospheric physics into vascular biomechanics, this topology-based framework yields a unified mechanistic language for describing near-wall hemodynamics, resolving blood flow complexity into interpretable and reproducible coherent fluid structures, enabling standardized hemodynamic phenotyping, and supporting hypothesis-driven studies of aneurysms and other cardiovascular diseases where greater fluid-mechanical specificity and interpretability may strengthen links between mechanobiology and clinical risk.

## INTRODUCTION

In external flows such as in the atmosphere, vorticity can be amplified by thermodynamic gradients to produce tornadic phenomena, whose destructive potential arises from intense “ground effects”. Vortical structures also develop in internal flows, including the cardiovascular system, where anatomy and cardiac pulsatility drive vorticity transport and blood flow interaction with the vessel wall. For decades, low wall shear stress (WSS), characteristic of slowly recirculating flow near bifurcations and bends (1), has been recognized as a “flow disturbance”, i.e., a modifier of blood– wall interactions (2). In combination with WSS multidirectionality, it can promote endothelial inflammation and atherosclerotic lesion development (*3*, *4*). Similarly, in aneurysms flow disturbances have been linked to rupture risk (*5*, *6*). In cerebral vessels, ruptured aneurysms may also exhibit more complex, unstable hemodynamics, richer intrasaccular vorticity, and elevated WSS gradients (*7*).

Despite decades of investigation, “flow disturbances” remain difficult to define, even though they underly the broader hemodynamic risk hypothesis of cardiovascular disease (*1*). Classical fluid mechanics describes disturbed flow through separation, stagnation, and recirculation. However, in the vasculature these patterns elude unambiguous classifications, and their biological implications depend strongly on context and vascular bed. This ambiguity has contributed to ongoing challenges in disentangling complex-but-benign physiological flow from deleterious hemodynamic patterns.

To address this gap, we recently identified near-wall vorticity kinematics as a key descriptor of the intravascular fluid structures that shape WSS topology (*8*, *9*). The present work leverages this framework to analyze cerebral aneurysm hemodynamics through coupled vorticity and WSS perspectives, interpreting the resulting near-wall fluid structures through a meteorological analogy. Specifically, we focus primarily on two damage-producing atmospheric events: tornadoes, rotating air columns coupled to the ground, and downbursts, descending nonrotating air columns that generate intense horizontal winds upon impact. We also consider multiple occurrences of these events taking place simultaneously and interacting along a developing line of storms.

As summarized in Fig. 1, combining fluid-mechanics theory with high-fidelity computational fluid dynamics (CFD)—the latter widely regarded as a credible tool for mechanistic investigation, relative flow-pattern analysis, and generation of testable hypotheses regarding the distribution of the hemodynamic stimuli imparted to the aneurysm wall (*10*)—we show that topological analysis of WSS and surface vorticity (SV) identifies near-wall tornadic fluid structures in cerebral aneurysms that are analogous to tornadic phenomena^1^. The analogy emphasizes potentially damaging near-surface actions at the atmospheric ground and at the vascular luminal surface. Time-resolved flow magnetic resonance imaging (4D flow MRI) acquisitions in patients presenting with cerebral aneurysm confirm that tornadic flow can also be observed, independently, *in vivo*.

**Fig. 1.**
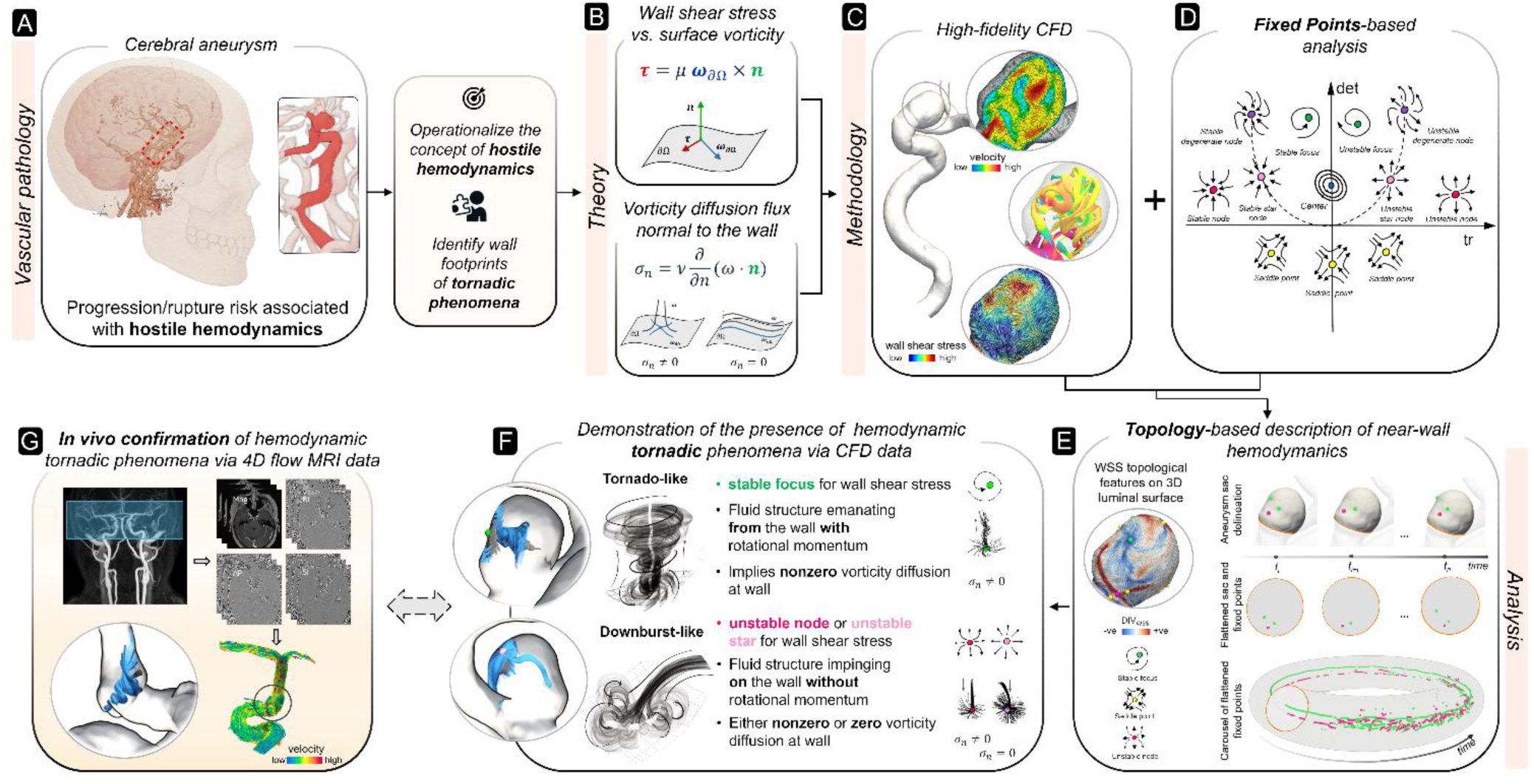
Overview of the study design and working hypothesis. (A) Cerebral aneurysm hemodynamics may influence aneurysm progression and rupture risk through hostile conditions, which are interpreted here through the near-wall signatures of tornadic-like phenomena. (B) The theoretical framework relates wall shear stress (WSS) to surface vorticity (SV) and examines wall-normal vorticity flux to define a topology-based taxonomy of near-wall fluid structures. (C) High-fidelity CFD is used to resolve patient-specific aneurysm hemodynamics. (D) Topological analysis identifies and classifies WSS and SV fixed points, including foci, nodes, saddles, and centers. (E) Topology-based analysis characterizes near-wall flow organization. Normalized WSS divergence (DIV_WSS_) is used to identify stable and unstable manifolds, while the fixed-point carousel method depicts the kinematics of WSS fixed points over the cardiac cycle. (F) CFD reveals tornadic hemodynamic phenomena and distinguishes rotational (tornado-like) from nonrotational (downburst-like) near-wall fluid structures based on WSS and vorticity topology. (G) 4D flow MRI provides independent evidence of tornadic-like intravascular flow structures *in vivo*. Illustrations in panels A and F were drawn by author T.N.

Per Fig. 1, the present study integrates prior work in cerebral aneurysm fluid mechanics, dynamical systems theory, and the mechanistic coupling between near-wall vorticity, WSS topology, and tornadic-like flow organization. These foundations are reviewed briefly below to motivate the analytical framework used in the Results and Discussion.

### Cerebral Aneurysms and their Flow Complexity

Cerebral aneurysms (Fig. 1A) are saccular outpouchings of brain arteries that occur in about 1 in 30 adults (*11*). Although the annual rupture risk is ∼1%, rupture commonly causes death or permanent disability (*12*). Increasing incidental detection of unruptured aneurysms by noninvasive imaging has sharpened the clinical dilemma because prophylactic treatment risks may approach rupture risk (*13*).

Aneurysm natural history reflects coupled mechanics and biology. Anatomically-driven flow disturbances impose focal mechanical stimuli on a vasculature already shaped by genetic susceptibility and environmental modifiers (*14*). Across initiation, growth, and rupture, hemodynamics, especially shear forces, regulates endothelial behavior through mechanotransduction and downstream signaling (*15*). Under physiological conditions the endothelium supports vascular homeostasis through antithrombotic and anti-inflammatory mediators, whereas altered hemodynamics can promote dysfunction and inflammation (*16*, *17*). The predilection of aneurysms for bifurcations and sharp bends, where flow complexity is intrinsically elevated, supports a causal contribution of local hemodynamics. Although rupture is ultimately a structural failure, wall degradation and vulnerability remain difficult to assess clinically; consequently, fluid mechanics is often used as a pragmatic proxy.

Because aneurysms are located deep within the brain, *in vivo* flow characterization was historically limited. Patient-specific CFD became feasible with clinical 3D x-ray angiography and affordable computing (*18*), and subsequent retrospective studies linked rupture status to both low (*5*) and high (*19*) WSS magnitude. CFD has been accepted by the clinical research community as an important tool for probing hemodynamic contributions to pathogenesis (10). More recent 4D flow MRI studies have enabled in vivo visualization of vortical structures, although current spatial and temporal resolutions still limit robust estimation of WSS and flow instability (20).

These complementary approaches *in silico* and *in vivo* have clarified many features of aneurysm flow while also revealing the incompleteness of current hemodynamic description. The low-versus-high WSS magnitude debate reflects a broader limitation of simpler biomechanical descriptors. The dual-pathway hypothesis (*21*) helped reconcile this partially by proposing that low, oscillatory WSS promotes an inflammatory-cell-mediated route to growth and rupture, whereas high WSS magnitude with steep spatial gradients promotes mural-cell-mediated wall degradation and thinning. Yet this dichotomy does not fully capture the vectorial, multidirectional nature of WSS, the spatiotemporal complexity of the mechanical action applied to the luminal surface, or its mechanistic coupling to near-wall flow organization.

Attention has therefore shifted from WSS magnitude alone to the intravascular structures that shape it. Vorticity has emerged as a marker of aneurysmal flow complexity, and higher complexity— quantified by vorticity and reflected by intrasaccular vortex corelines—has been associated with increased rupture risk (*7*). Because vorticity also governs convective mixing and near-wall mass transport, its local organization may influence the accumulation of inflammatory or atherogenic species and oxygen delivery to the wall. Impaired transport in the near-wall domain may upregulate metalloproteinases, promote extracellular matrix degradation, weaken the wall, and favor aneurysm formation and remodeling (*16*). Together, these observations motivate this study: a fluid-mechanics framework that links near-wall fluid structures, WSS topology, and biologically relevant wall exposure.

### Taxonomy of Near-Wall Fluid Structures via Fixed-Point Analysis

Topological analysis of vectorial quantities provides a compact, geometrically invariant description of their organization. In dynamical systems terms, a topological skeleton is defined by fixed points and manifolds connecting them. Fixed points are locations where the field vanishes; manifolds indicate directions along which nearby field lines are attracted or repelled. Depending on local field-line geometry, fixed points are classified as saddles, centers, foci, and nodes, with foci and nodes further divided into stable and unstable types (Fig. 1D). Saddles attract and repel along distinct directions, centers are encircled by closed orbits, foci by spiral trajectories, and nodes by converging or diverging field lines, including degenerate or star-node configurations.

Building on the theoretical correspondence between topological skeletons of WSS and SV, we previously developed a physically grounded taxonomy of admissible fixed-point pairings based on type, stability, and their mechanistic association with specific near-wall fluid structures (*9*). The framework further distinguishes these structures by the local presence of wall-normal vorticity diffusion flux (Fig. 1F), thereby resolving differences in near-wall vorticity kinematics (*9*). This provides a principled way to identify which intravascular fluid structures are most likely to imprint on the vessel wall. It also offers compact language for connecting detailed flow fields to specific classes of biomechanical action on the vessel wall.

### Fluid Mechanics of Tornadic Phenomena

We use the topology-based taxonomy to show that cerebral aneurysm hemodynamics exhibit near-wall configurations analogous to several atmospheric events: tornado-like columnar vortices connected to the aneurysmal wall, downburst-like vertical fluid drafts that impinge on the wall without rotation (Fig. 1F, Fig. 2A, B). In the atmosphere, tornadoes develop when pre-existing vorticity is tilted, stretched, or aggregated by vertical drafts in a sheared environment. Buoyant updrafts can rotate a horizontal vortex tube toward the vertical; convergence toward the resulting low-pressure core then amplifies tangential speed by angular-momentum conservation, narrowing and intensifying the vortex. Near the surface, friction and convergence further strengthen the circulation and anchor it to the ground.

**Fig. 2.**
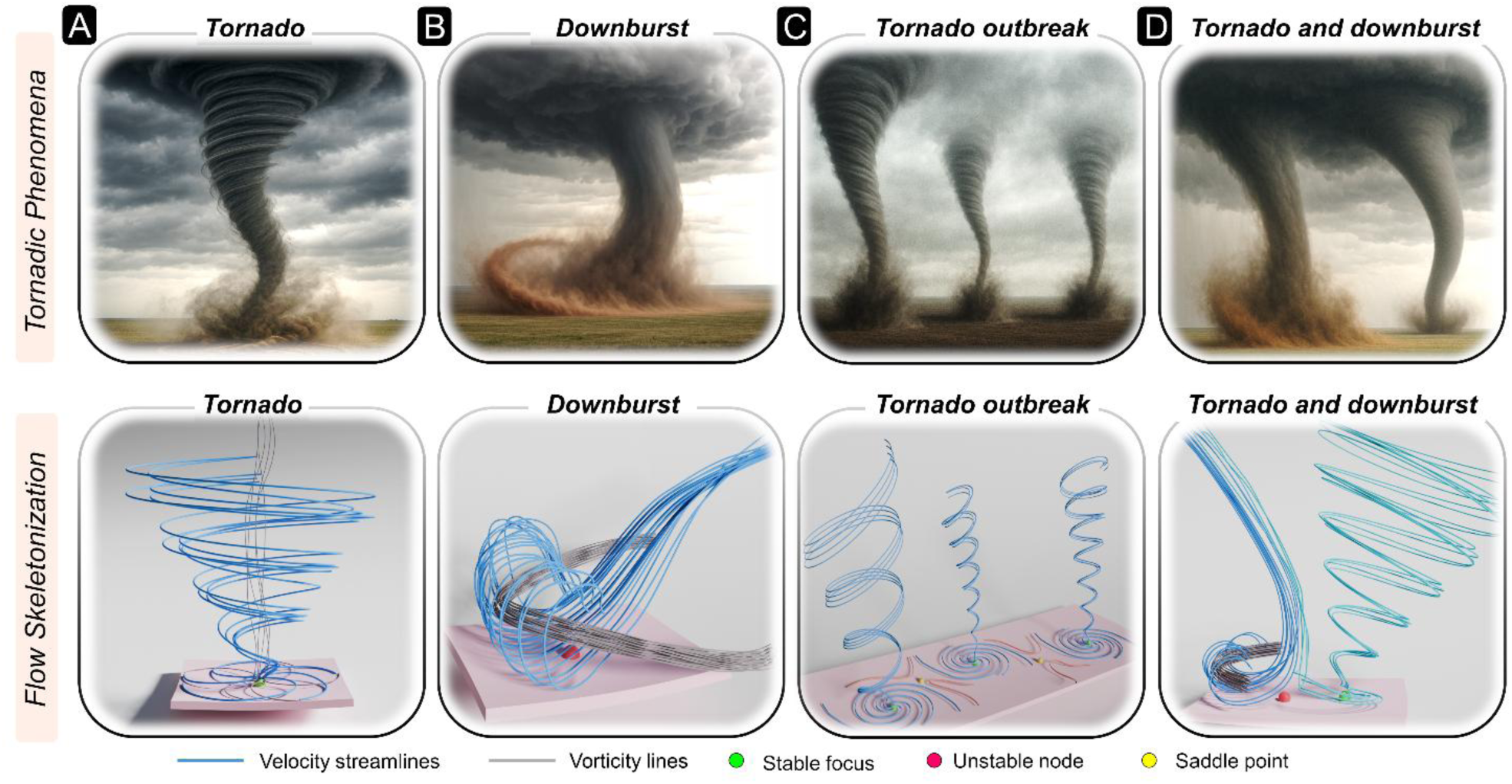
Tornadic phenomena and their flow skeletonization. Atmospheric tornadic phenomena are illustrated* in the upper row and represented by idealized fluid-mechanical skeletons in the lower row. Skeletons are defined by velocity streamlines, shown in light blue, and vorticity lines, shown in gray. (A) Tornado-like structure. A vertical rotational columnar flow is represented by velocity streamlines that wrap spirally around nearly wall-normal vorticity lines and connect to a WSS stable focus, marked by the green sphere, at the wall. (B) Downburst-like structure. A nonrotational descending columnar flow spreads laterally after wall impact, producing near-wall deflection of streamlines and a wall-parallel vortex, visualized by streamlines wrapping around horizontal vorticity lines. This topology connects to a WSS unstable node, marked by the magenta sphere. (C) Tornado-outbreak-like structure. Multiple tornado-like structures form in proximity, producing a WSS skeleton with adjacent WSS stable foci, marked by green spheres, separated by saddle points, marked by yellow spheres. (D) Coupled tornado-downburst-like structure. Rotational and nonrotational columns coexist, connecting respectively to a WSS stable focus, marked by the green sphere, and to a WSS unstable node, marked by the magenta sphere. *The lower-row skeletonizations were created using the open-source Blender 3D graphics suite. The upper-row illustrations were generated using ChatGPT (OpenAI, Inc.) to provide copyright-free atmospheric renderings that visually parallel the corresponding fluid-mechanical features. The authors take full responsibility for the conceptual and scientific accuracy of the synthetic illustrations.

The damage potential of a tornado arises from the near-surface winds within the rotating column rather than from the visible condensation funnel, which may not extend to the ground or may be absent altogether. Although velocity vanishes exactly at the boundary because of the no-slip condition, large tangential and convergent motions can occur immediately above it. Multiple tornadoes may also occur within the same large-scale severe-weather setting, producing distinct but overlapping damage tracks; such multi-tornado systems motivate our consideration of compound near-wall structures in aneurysms (Fig. 2C).

Convective storms can also generate damaging near-surface flow without a rotating vertical core. The canonical example is the downburst (*22*), where rapidly descending air impacts the ground and spreads outward, creating intense divergent horizontal winds rather than the convergent rotating pattern of a tornado. Tornadoes and downbursts can coexist or interact, yielding hybrid wind fields that complicate interpretation of near-surface damage. Tornado–downburst combinations show how rotational and non-rotational near-surface actions can coexist within a single event (Fig. 2D).

Although less directly comparable to tornadoes or downbursts, roll clouds—elongated tube-like structures that rotate about a horizontal axis roughly parallel to the ground—illustrate coherent rotation without a vertical core and can coincide with strong horizontal winds. Their inclusion broadens the range of atmospheric analogues used to interpret near-wall hemodynamic organization.

More generally, tornadic phenomena can be interpreted by skeletonizing the flow field using vortex lines and velocity streamlines. Intertwined vortex lines and streamlines indicate strongly rotational, coherent structures, whereas non-intertwined patterns indicate flow dominated by translation and viscous effects. Fig. 2 summarizes these archetypes and their skeletonized representations, which provide the physically grounded basis for interpreting the hemodynamic phenomena reported in the following section.

## RESULTS

Before drawing parallels between atmospheric events and hemodynamic phenomena in cerebral aneurysms, it is essential first to appreciate the inherent complexity of intra-aneurysmal blood flow and the need for advanced modeling and visualization techniques capable of capturing a broad spectrum of hemodynamic behaviors associated with disease. Movies 1-3 illustrate three distinct aneurysmal flow phenotypes over the cardiac cycle, from stable to highly unstable, using instantaneous velocity magnitude, vortical structures identified by swirling strength (a measure of amount of local fluid rotation; Methods), and the WSS topological skeleton. These data derive from high-fidelity CFD simulations of three anatomically realistic cases (Fig. 3A) representing stable (Case S; Movie 1), moderately unstable (Case Um; Movie 2), and highly unstable (Case Uh; Movie 3) flow regimes (*23*). Intra-aneurysmal flow leaves a measurable imprint on the vessel wall through spatially and temporally varying shear forces that reflect the underlying hemodynamic complexity. These mechanical cues are captured by spatiotemporal changes in the WSS topological skeleton, obtained by globally mapping fixed points and manifolds—the latter identified from the divergence of normalized WSS, as described in the Methods. As illustrated in Fig. 1E and Movies 1-3, the analysis focuses on WSS stable foci and WSS unstable nodes to identify tornadic phenomena.

**Fig 3.**
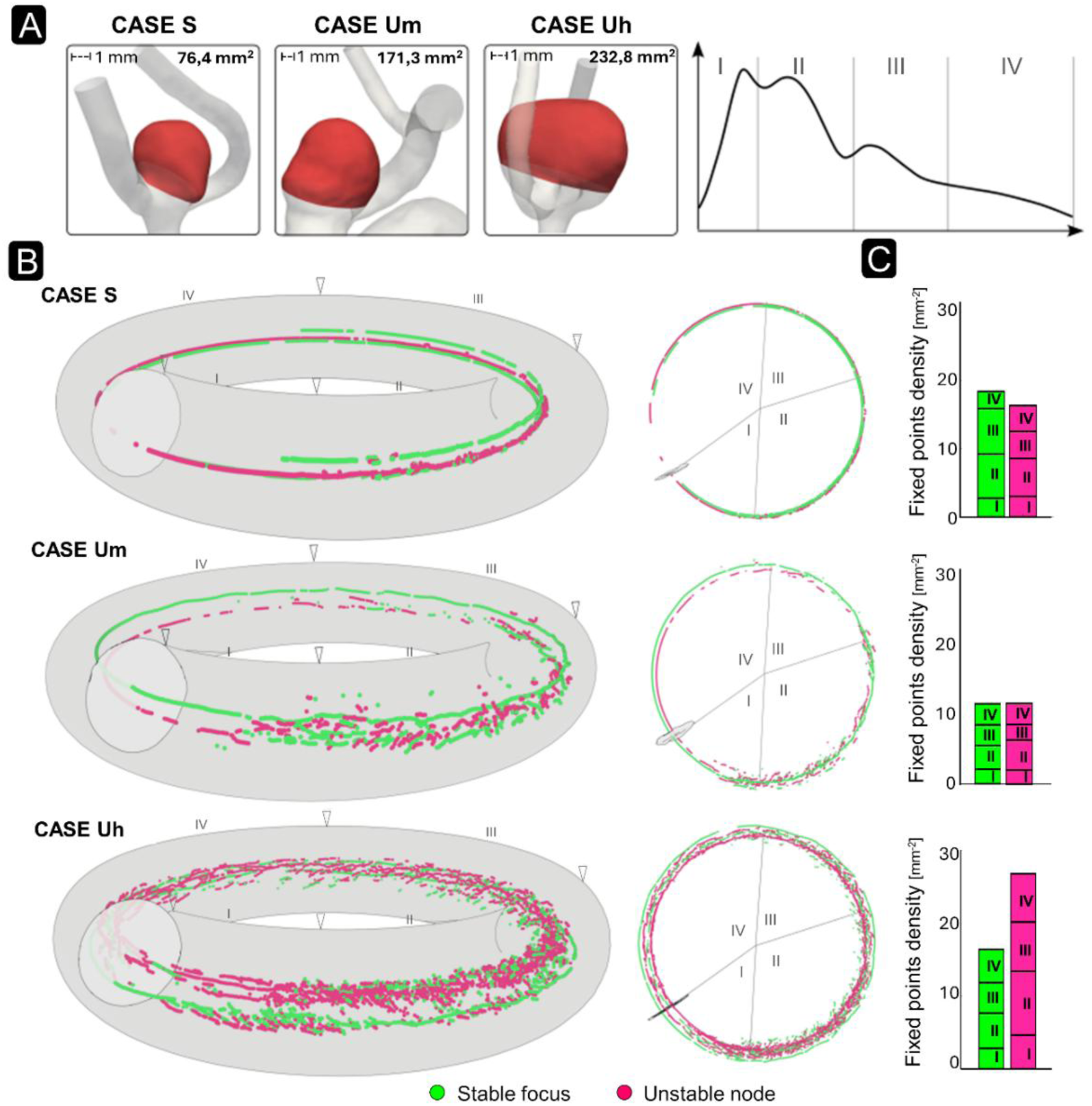
High-fidelity CFD models and fixed-point carousel visualization. (A) Reconstructed geometries of three representative unruptured cerebral aneurysms: Case S, with predominantly stable flow; Case Um, with moderately unstable flow; and Case Uh, with highly unstable flow. Aneurysm sacs are shown in red, with scale bars for reference, and their luminal surface area are also reported. The inflow waveform shows the prescribed volumetric flow-rate profile over the cardiac cycle, divided into four characteristic phases, I–IV. (B) Fixed-point carousel visualization of the spatiotemporal evolution of WSS fixed points theoretically associated with tornadic flow structures. WSS stable foci, marked by green spheres, are associated with rotational tornado-like structures, whereas WSS unstable nodes, marked by magenta spheres, are associated with nonrotational downburst-like structures. In each carousel, the aneurysm sac is flattened onto a parametric surface and swept through time, enabling joint visualization of fixed-point location and cardiac-cycle timing. The same carousel is shown from an oblique view and from above to highlight temporal transitions and spatial clustering of near-wall topological features. (C) Luminal surface density of each fixed-point type, calculated as the cumulative number of fixed points over each of the four characteristic phases (I-IV), and normalized by the respective case’s sac surface area.

The intra-aneurysmal blood flow complexity revealed by CFD is corroborated by *in vivo* 4D flow MRI from two representative patients (Cases M1 and M2; Movie 4). These exhibit blood flow dynamics within the aneurysm characterized by the presence of tornadic fluid structures. Therefore, they demonstrate that sufficiently large tornadic-like fluid structures are directly observable *in vivo*, under standard clinical acquisition conditions.

Aneurysmal flow complexity is further illustrated in Fig. 3B, which shows the kinematics of WSS fixed points across the cardiac cycle using the fixed-point carousel methodology recently introduced in (*23*) and described in the Methods. The kinematics of stable foci and unstable nodes— WSS fixed points theoretically associated with tornadic phenomena—reveal distinct near-wall flow phenotypes across the three cases. Case Uh (highly unstable) exhibits substantially more WSS fixed points than Case Um (moderately unstable) and Case S (stable), even after accounting for its larger surface area. Across all cases, the carousels reveal numerous nearly stationary fixed points, indicating luminal locations occupied persistently by WSS fixed points throughout the cardiac cycle. In Case Uh, however, most fixed points appear and disappear intermittently—particularly during the post-systolic deceleration (phase II)—reflecting the elevated hemodynamic instability. Conversely, Case S exhibits a sparsely populated carousel, with small numbers of stable foci and unstable nodes remaining nearly stationary for most of the cardiac cycle, consistent with a stable flow regime. Case Um exhibits intermediate fixed-point kinematics, supporting the ability of the carousel to summarize gradations of hemodynamic complexity. In general, the incidence of fixed points, and of the associated near-wall tornadic fluid structures, rises after peak systole, as flow decelerates, consistent with the expected onset of flow instabilities with adverse pressure gradients and competing currents within the aneurysm sac (*24*, *25*). The surface densities of stable foci and unstable nodes (associated with tornado- and downburst-like fluid structures, respectively) differ across cases, with Case Uh exhibiting higher density of unstable nodes (Fig. 3C), indicating a higher density of downburst-like structures. Notably, these counts do not account for the relative strength or persistence of individual tornadic fluid structures, a point that will be addressed further on.

As an illustrative example, Fig. 4A shows tornado-like fluid structures in the aneurysmal near-wall domain of Case Um. Using the fixed-point carousel, we localized in space and time a WSS stable focus, whose presence implies a nonzero wall-normal vorticity diffusion flux—a topological configuration invariably associated with a rotational fluid structure emanating from the wall (Fig. 1F) (*9*). In the near-wall region surrounding the WSS stable focus, which corresponds to a stable SV focus: velocity streamlines wrap around nearly straight vortex lines, consistent with the existence of a rotating fluid column that expands toward the core of the aneurysmal sac (Fig. 4A, right panel), thus revealing the characteristic skeleton of a tornado-like fluid structure as shown in Fig. 2A. A similar tornado-like fluid structure was independently observed *in vivo* using 4D flow MRI (Fig. 4B). Despite the limited spatial resolution of phase-velocity data, the *in vivo* WSS topology confirms that the tornado-like fluid structure observed in Case M1 coincides with a WSS stable focus configuration, as dictated by theory (*9*).

**Fig 4.**
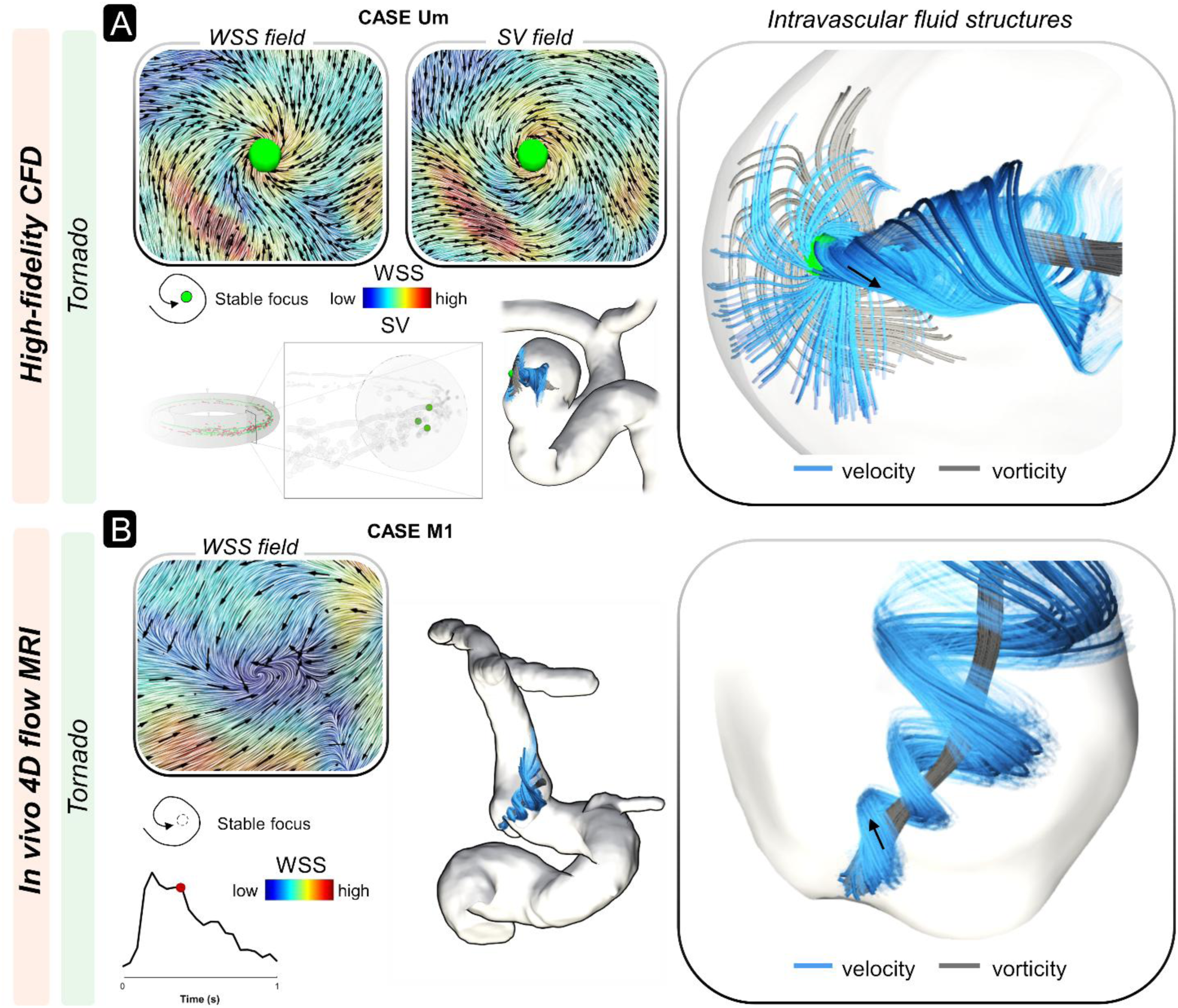
Representative tornado-like fluid structures in cerebral aneurysms. (A) High-fidelity CFD of Case Um. WSS and SV are shown on the aneurysm wall as unit-vector fields to emphasize their local topology. Both fields exhibit a stable focus, marked by the green sphere, consistent with a locally nonzero wall-normal vorticity flux. The WSS fixed-point carousel indicates three WSS stable foci at the analyzed time point; one focus is shown in detail. Aneurysmal velocity streamlines (blue) wrap helically around vorticity lines (gray), forming a tornado-like fluid structure anchored at the stable focus, as predicted by theory. (B) *In vivo* 4D flow MRI in Case M1 shows an analogous topology. WSS topology is visualized using unit vectors and line-integral convolution. An inferred WSS stable focus is associated with helically organized velocity streamlines around vorticity lines extending into the aneurysm sac, consistent with a tornado-like structure. Black arrows indicate the local near-wall flow direction.

Applying the same framework, the fixed-point carousel was employed to localize the presence of downburst-like fluid structures, which are associated with WSS unstable nodes or stars. According to the underlying theory, the presence of such nodes necessarily coexists with a nonrotational fluid structure impinging on the wall (Fig. 1F) (*9*). Using Case Um and its WSS fixed-point carousel as an illustrative example, Fig. 5A shows that in the near-wall region around the identified WSS star node—which in this case corresponds to a center SV fixed point—instantaneous velocity streamlines and vortex lines delineate a downburst-like fluid structure. Consistent with the schematic in Fig. 2B, velocity streamlines remain nearly straight, indicating a nonrotational column of fluid that impacts the wall; upon impingement, streamlines wrap around vortex lines, indicating the formation of a vortex structure with its axis oriented parallel to the wall. As with the tornado-like, downburst-like fluid structures were also independently observed *in vivo* using 4D flow MRI (Fig. 5B). For the illustrative Case M2, the estimated *in vivo* WSS topology confirms that the identified downburst-like fluid structure coincides with an unstable WSS node configuration, as dictated by theory (*9*).

**Fig 5.**
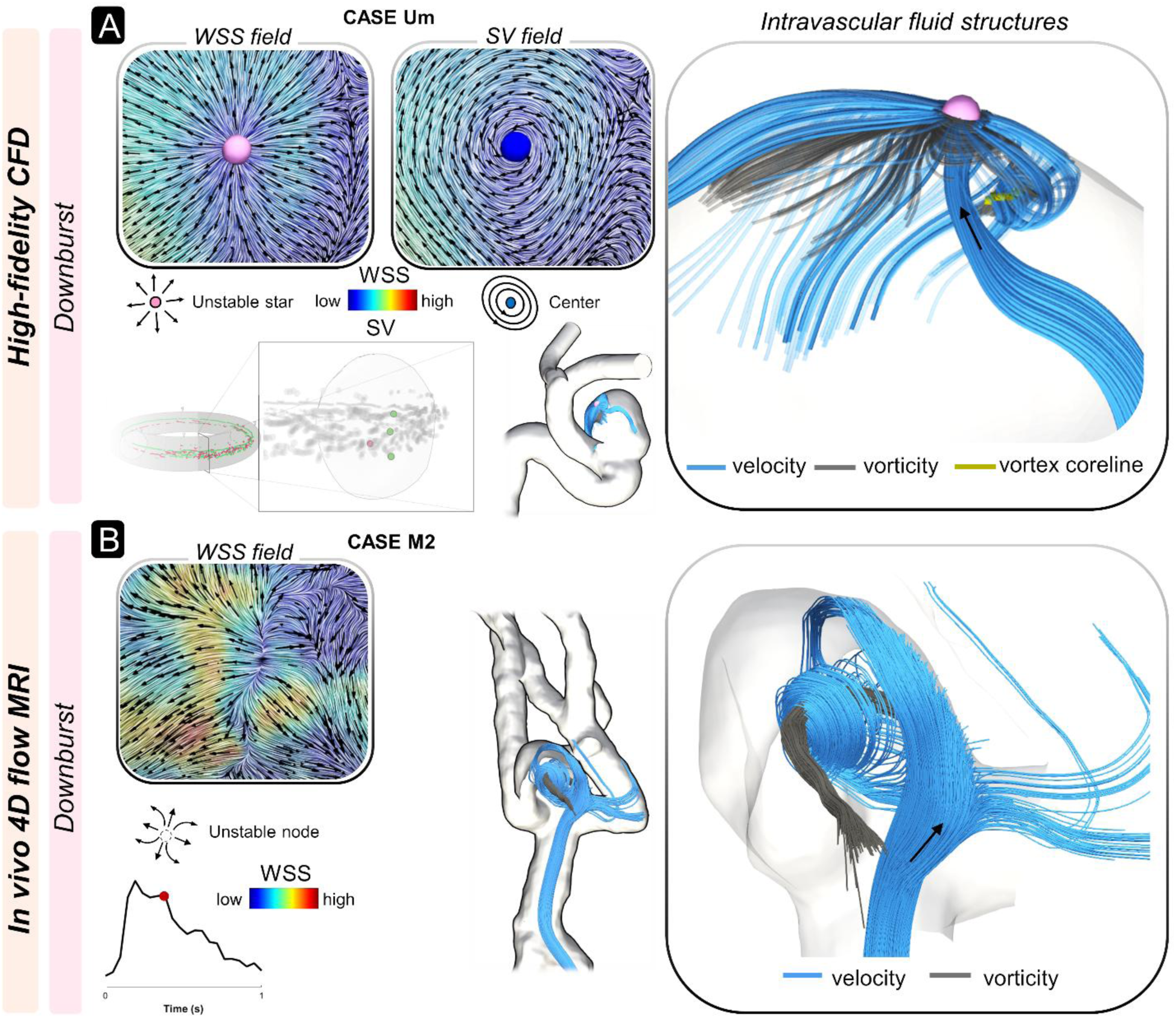
Representative downburst-like fluid structures in cerebral aneurysms. (A) High-fidelity CFD of Case Um. WSS and SV are shown on the aneurysm wall as unit-vector fields to emphasize their local topology. The WSS field contains an unstable star node, marked by the pink sphere, that coincides with a center in the SV field, marked by the blue sphere. This WSS–SV configuration is consistent with zero wall-normal vorticity flux, as predicted by theory. The WSS fixed-point carousel localizes the event in space and time. Velocity streamlines (blue) connected to the fixed point and vorticity lines (gray) delineate a nonrotational jet-like structure impinging on the aneurysm wall. After impact, the streamlines wrap around a near-wall vortex coreline (yellow) indicating formation of a wall-parallel vortex structure. (B) *In vivo* 4D flow MRI in Case M2 shows an analogous topology. WSS topology is visualized using unit vectors and line-integral convolution. An inferred WSS unstable node is associated with a downburst-like structure. Black arrows indicate the local near-wall flow direction.

The WSS fixed-point carousels of Fig. 3B enabled identification of compound tornadic events analogous to the atmospheric archetypes in Fig. 2, including tornado outbreaks and tornado-downburst configurations. In CFD Case Um, for example, the simultaneous presence of three stable WSS foci corresponded to three nearby tornado-like fluid structures in the near-wall region (Fig. 6A), as dictated by theory and illustrated by the skeletonization in Fig. 2C. A tornado-outbreak-like configuration within the aneurysm sac requires the WSS topological skeleton to satisfy two conditions. First, it must obey the kinematic constraints on the admissible number and types of fixed points—as prescribed by Poincaré-Hopf theorem (*26*). Second, it must exhibit a spatial ordering in which stable foci alternate sequentially with saddle points. This WSS topological skeleton, which can be readily identified from the manifolds connecting fixed points (Fig. 6A, lower left panel) (*8*, *27*), is uniquely consistent with a tornado outbreak-like configuration in the near-wall domain. In CFD Case S, the WSS fixed-point carousel revealed a closely co-located pair consisting of a stable focus and an unstable node which, by topology, cannot be directly connected by a manifold (Fig. 6B). This configuration corresponded to a near-wall flow pattern analogous to a tornado adjacent to a downburst, as highlighted by the skeletonization in Fig. 2D. Consistent with theory, a rotational fluid column expanding toward the sac core maps to a stable WSS focus, whereas a nonrotational, wall-impinging fluid structure that generates a tangential near-wall current maps to a unstable WSS node. The corresponding SV fixed points displayed the expected types and stability properties (see Methods).

**Fig 6.**
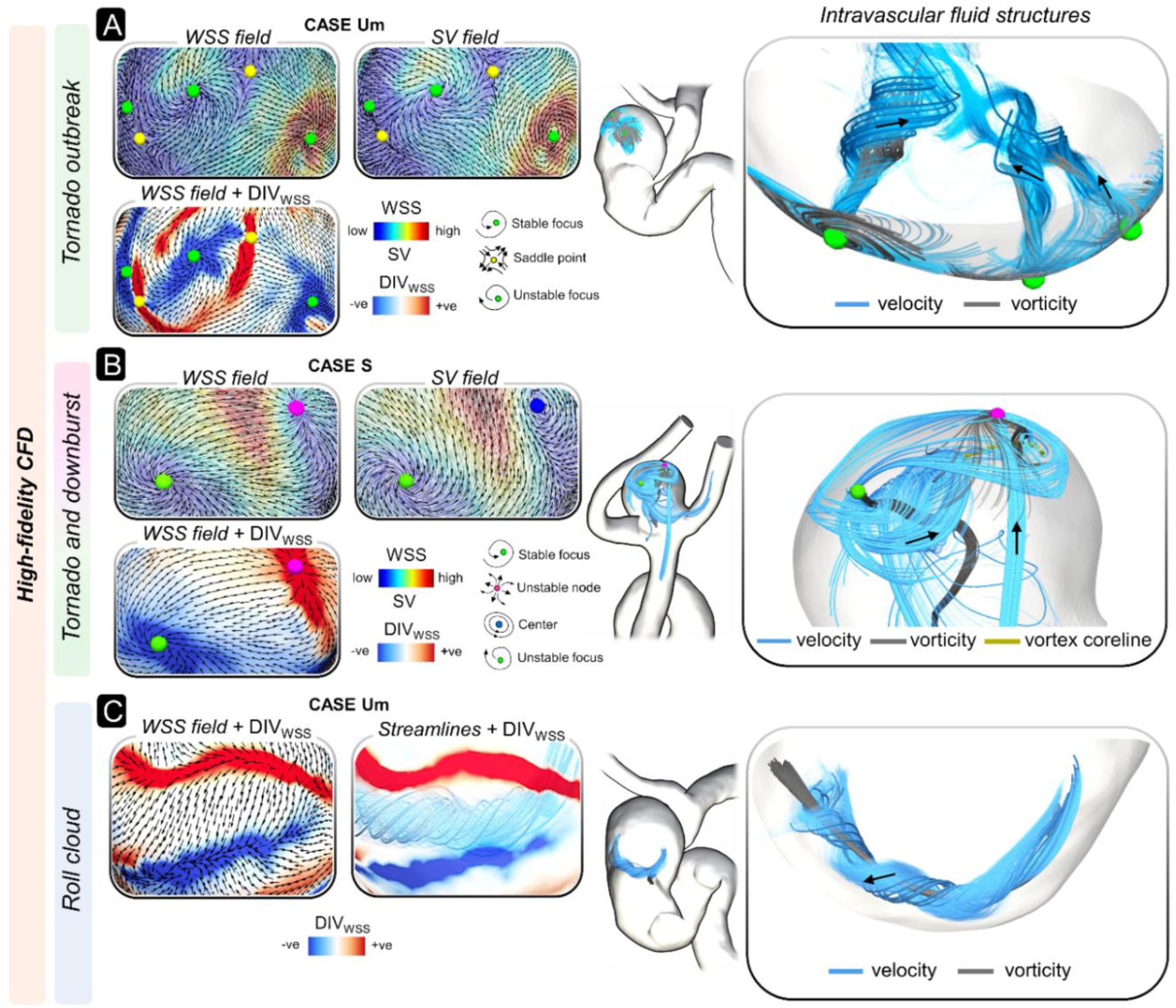
Representative combinations of other tornadic phenomena in cerebral aneurysms. WSS and SV are shown on the aneurysm wall as unit-vector fields to emphasize their local topology. Normalized WSS divergence is used to identify stable and unstable WSS manifolds. (A) Tornado-outbreak-like topology in Case Um. Three simultaneous WSS stable foci, marked by green spheres, each coincide with SV stable foci, identifying multiple tornado-like structures. These foci alternate with WSS saddle points, marked by yellow spheres. Unstable WSS manifolds, identified by negative (-ve) DIV_WSS_ values, connect saddle points to adjacent stable foci. Velocity streamlines (blue) associated with the WSS stable foci and vorticity lines (gray) confirm multiple coexisting tornado-like structures, consistent with an outbreak topology. (B) Coupled tornado–downburst-like topology in Case S. A WSS stable focus, marked by the green sphere and corresponding to a SV unstable focus, coexists with a WSS unstable node, marked by the magenta sphere, and corresponding to a SV center, marked by the blue sphere. This configuration identifies adjacent rotational tornado-like and nonrotational downburst-like structures. Velocity streamlines and vorticity lines confirm the combined near-wall flow system. (C) Roll-cloud-like near-wall vortex in Case Um. Although not part of the fixed-point taxonomy, this structure is revealed by the WSS manifold pattern: alternating stable and unstable WSS manifolds, identified by positive (+ve) and negative (-ve) DIV_WSS_ values, bound a tangentially rolling tubular vortex along the aneurysm wall. Black arrows indicate the local near-wall flow direction.

We also examined tornadic phenomena not directly anchored to aneurysmal wall, including roll-cloud-like fluid structures that may nonetheless produce a “ground effect”. Roll clouds are relevant atmospheric analogues because they are tube-shaped vortex structures rotating about an axis parallel to the ground (*28*, *29*), and they resemble funnel cloud-like structures reported within aneurysm sacs (*30*, *31*). Using swirling strength (see Methods) to visualize vortex structures, together with WSS fixed points and manifolds—identified using DIV_WSS_ maps (see Methods)—to delineate the WSS topological skeleton (Movies 1-3), we found that coherent tube-shaped vortices can also arise in near-wall regions that lack fixed points and are bounded by stable and unstable manifolds. These tubular vortices are topologically detached from fixed points and adopt a roll-cloud-like configuration, with their axis of rotation oriented parallel to the wall. This behavior is illustrated in CFD Case Um (Fig. 6C), where a tubular fluid structure rolls parallel to the luminal surface. The interpretation for the observed phenomenon is that nearby WSS manifolds delimit a region where near-wall currents interact or oppose each other, as suggested by the WSS map in Fig. 6C, used here as a surrogate of the near-wall flow field (see eq. (1); Methods). This interaction may induce fluid lift; in the presence of a wall-normal shear gradient, the lifted fluid curls into a tube rotating about an axis parallel to the luminal surface, resembling a classical shear-driven roll atmospheric cloud.

## DISCUSSION

The findings of this study indicate that cerebral aneurysm hemodynamics can be conceptualized within a topological framework that parallels atmospheric tornadic phenomena. This cross-disciplinary analogy extends the organizing principles and terminology from meteorological fluid mechanics to vascular biomechanics, adopting a common descriptive language for distinct atmospheric events and near-wall blood flow patterns. Viewed through this lens of observation, the topological approach offers a means to disentangle the intrinsic complexity of aneurysmal hemodynamics—and benign hemodynamic complexity in other cardiovascular settings—from superimposed flow disturbances that may accompany pathological vessel wall remodeling and adverse clinical outcomes. In this study, we focused on cerebral aneurysms as a representative vascular setting in which highly complex hemodynamics makes biologically and clinically adverse flow patterns difficult to identify. In this context, conventional fluid-mechanical descriptors often provide limited support for stratifying the risk of aneurysm growth or rupture. We therefore tested the proposed topological framework in this particularly challenging pathological scenario, where greater fluid-mechanical specificity and interpretability may be especially valuable.

The proposed analogy is intended strictly at the level of flow organization, particularly vorticity kinematics in the near-wall domain. Given the theoretical correspondence between SV and WSS topology (*9*), and the established role of WSS in vascular pathophysiology (*17*, *32–34*), the “ground effects” of atmospheric tornadic systems motivate the hypothesis that analogous confined near-wall fluid structures may adversely affect the luminal surface of a vessel. Within this framework, the tornadic aneurysmal fluid structures can be recast as distinct hemodynamic actions with potentially distinct mechanobiological implications. Tornado-like fluid structures, via sustained rotational shear, may be associated with chronic endothelial stimulation and inflammation (*16*), whereas the concurrent emergence of multiple such structures in a tornado-outbreak configuration may extend these effects over larger regions of the aneurysmal sac. Downburst-like fluid structures may produce localized wall impingement and tangential outflow, imposing impulsive mechanical loading and cyclic expansion–contraction actions on the luminal surface, analogous to the damaging near-ground horizontal winds of atmospheric downbursts. Simultaneous coexistence of tornado- and downburst-like configurations within close spatial proximity may further intensify near-wall hemodynamic gradients that nonphysiologically amplify mechanical stimuli to the vessel wall. Finally, roll-cloud-like fluid structures —persistent tangential vortices sustained by shear gradients and modulated by pulsatility—may likewise impose expansion-contraction forces on the wall. Collectively, these tornadic configurations delineate a spectrum of coherent near-wall flow phenomena capable of generating both chronic and acute mechanical stresses that may contribute to aneurysm progression and rupture.

Beyond the analogy, the topological framework provides a mechanistic taxonomy of near-wall fluid structures. It offers a principled means of consolidating velocity streamlines, vortex lines, and WSS fixed points and manifolds into a unified description. These flow features have often been used separately and/or interpreted in a broader descriptive fluid-mechanical sense (*35–38*). As geometric tracers of fluid motion, they delineate coherent flow patterns and connect three-dimensional near-wall flow organization to the two-dimensional WSS topology on the luminal surface, thereby enabling a physical interpretation of complex aneurysmal hemodynamics.

Such a theory-driven, topology-based taxonomy provides a reproducible framework for classifying vascular flow patterns, moving beyond purely descriptive and relatively non-specific in silico or *in vivo* observations, and enabling mechanistically grounded interpretation of cerebrovascular hemodynamics. By offering a quantitative and geometrically invariant description of near-wall hemodynamics organization, the proposed framework is useful not only for identifying near-wall fluid structures, but also for interpreting and, when appropriate, classifying them. In this way, it may help define and categorize blood-flow patterns that could exert adverse biological effects on the vessel wall, thereby addressing the still ambiguous notion of “disturbed flow”. By contrast, conventional hemodynamic quantities have often yielded conflicting conclusions (*21*). These quantities have also been derisively termed “confounding factor dissemination” (*39*) and “an alphabet soup” (*40*), because they capture only selected aspects of the inherently multifaceted WSS field (*41*, *42*), or use vorticity as a global measure of flow complexity without distinguishing near-wall contributions from vorticity transport in the bulk flow (*30*). The topology-based approach, instead, identifies fixed points and manifolds that partition near-wall flow into regions of convergence, divergence, and rotation. In doing so it delineates the structural skeleton of the near-wall flow field and enables a mechanistic characterization of flow disturbances via specific topological configurations, rather than through loosely defined descriptions such as stagnation, separation, impingement, or recirculation.

To illustrate the potential benefits of our approach, Fig. 7A maps the accumulated residence times of WSS fixed points associated with tornadic phenomena (stable foci and unstable nodes), weighted by their nominal strength (*RT∇_xfp_*; see Methods). These maps reveal distinct focal hot spots of tornado- and downburst-like ground effects, whereas conventional WSS-derived quantities commonly associated with rupture risk (TAWSS and OSI (*5*, *43*), introduced in the Supplementary Materials, Methods) display multiple, often spatially extended regions of extreme values (Fig. 7B). For example, in Case S, both tornado- and downburst-associated hot spots for RT∇*_xfp_* coincide with regions of elevated OSI, despite their potentially different mechanobiological implications. In the same Case S, broad luminal surface areas of low and high TAWSS correspond, respectively, to comparatively localized hot spots of tornado- and downburst-like activity. In Cases Um and Uh, extended regions of low TAWSS and high OSI—often interpreted as rupture-associated signatures (15)—occur in areas without detectable tornadic activity. Notably, per Fig. 7C, the aneurysmal flow complexity quantified by vortex corelines (a previously proposed nominal predictor of rupture status (*30*); see Methods) may aggregate many vortex structures that are not anchored to the near-wall domain and thus may be less specific to mechanobiological activity at the wall.

**Fig 7.**
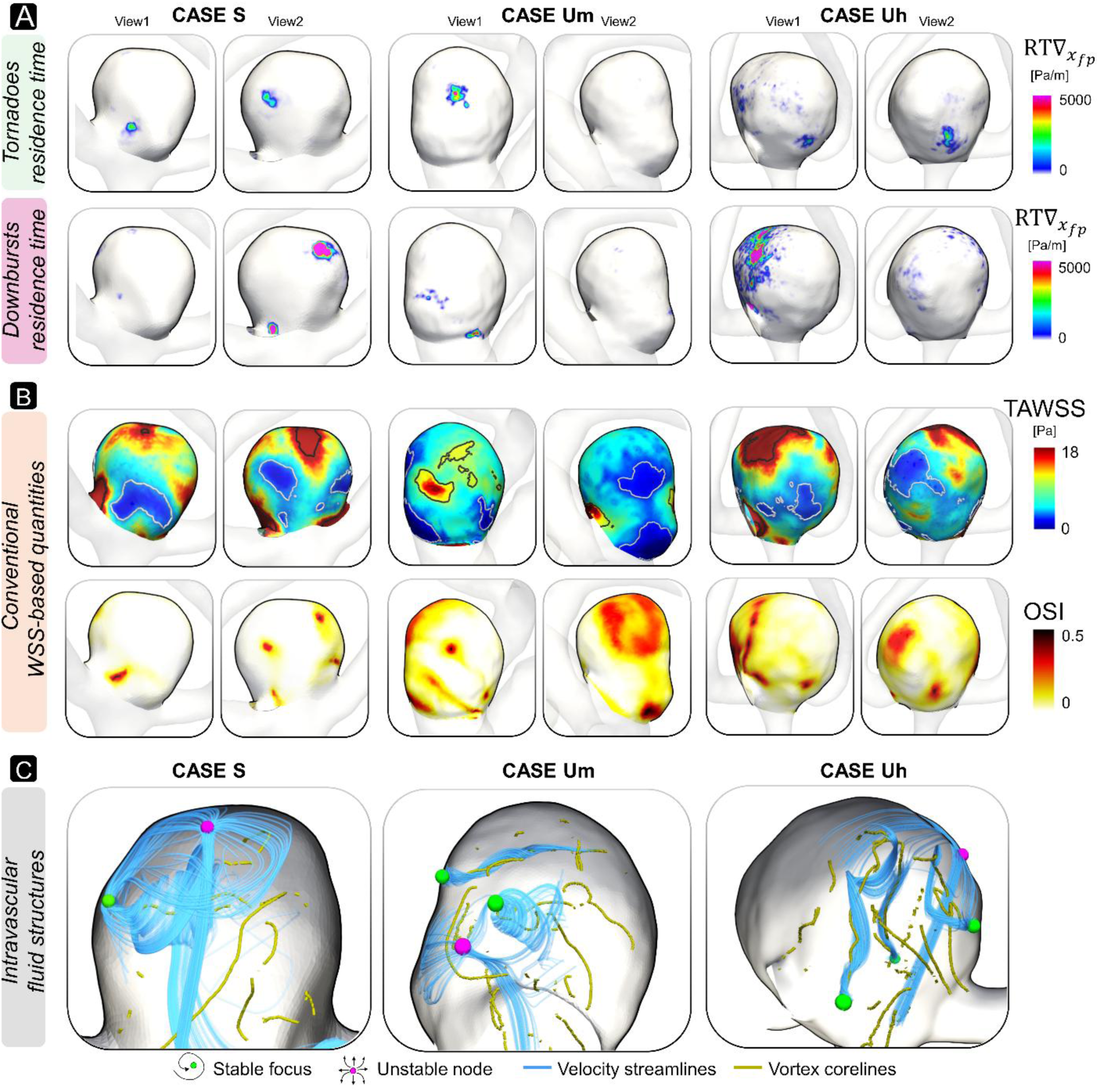
Near-wall and intravascular flow quantities in cerebral aneurysms. (A) Accumulated residence time of WSS fixed points (RT∇*_xfp_*) on the aneurysm sac over the cardiac cycle, weighted by the local magnitude of WSS divergence, which quantifies the intensity of local WSS contraction or expansion. Maps are shown separately for WSS stable foci, associated with tornado-like structures, and WSS unstable nodes, associated with downburst-like structures. These maps identify luminal regions persistently exposed to tornadic near-wall flow phenomena. Two views are shown for each case. (B) Conventional WSS-based quantities on the aneurysm sac: time-averaged WSS (TAWSS) and oscillatory shear index (OSI). Dark-gray and light-gray contours outline high- and low-TAWSS regions, respectively, defined by the 90^th^ and 10^th^ percentiles of the sac-wise TAWSS distribution. Two views are shown for each case. (C) Instantaneous intravascular flow organization within each aneurysm sac. Velocity streamlines seeded from WSS stable foci and WSS unstable nodes are shown in blue to visualize tornado-like and downburst-like structures, respectively, together with instantaneous vortex corelines shown in yellow.

Because the CFD cases analyzed here were drawn from unruptured aneurysms undergoing treatment to prevent rupture, we can only speculate that tornadic flow phenomena may provide more specific insight into focal risks of aneurysm growth or rupture, owing to their associated “ground effects”. Future investigations in patient cohorts will therefore be essential to elucidate the mechanobiological impact of near-wall vorticity dynamics—topologically analogous to those observed in atmospheric systems—on cerebral aneurysms formation, progression, and rupture, particularly through their relationship to the WSS topological skeleton. Prior CFD studies have already sought to relate hemodynamic extrema to focal wall abnormalities, including rupture sites (*38*), thin versus thick regions observed intraoperatively as proxies for wall composition or structural integrity (*44*), regions of irregular pulsation on dynamic computed tomography imaging (*45*) as proxies for wall weakness, and localized MRI contrast enhancement as a proxy for wall inflammation (*46*). Collectively, these observations point to the possibility that focal tornadic flow phenomena can be linked to measurable and more specific proxies of biological and biomechanical wall instability, thereby refining our understanding of the mechanobiology of wall degradation. Numerous CFD studies of large cohorts of ruptured and unruptured cerebral aneurysms have already been published (*10*), and could, in principle, be further leveraged through additional topological post-processing to relate the classifications introduced here to biomechanics-driven biological events, with the final goal of expanding the mechanobiological comprehension of the disease and improving clinical rupture risk stratification. Moreover, similarly extensive CFD studies of aneurysmal or dilated ascending and abdominal aortas—vascular pathologies that exhibit similar focal phenomena analogous to those of cerebral aneurysms—offer an opportunity to extend the proposed topological framework and assess its capability to distinguish physiological flow complexity from hostile hemodynamics with adverse mechanobiological consequences.

Finally, the concordance of theory with CFD simulations and the *in vivo* direct observation of tornadic phenomena with 4D flow MRI indicate translational potential. When 4D flow MRI adequately resolves the relevant scales of tornadic fluid structures, these can be observed and reconciled quantitatively with theoretical expectations. At current spatial resolutions (∼0.8-1.0 mm^3^ (*47*)), our study shows that tornadic structures can be readily observed in the aneurysm sac. Further identification and reconciliation of finer vortex structures and topological features may, however, require still-uncommon higher magnetic field strengths and/or physics-informed data augmentation strategies (*48*). Mechanobiological relevance is further suggested by prior work linking WSS topological skeleton features to biochemical transport at the blood-vessel interface (*27*, *49*, *50*), atherosclerotic lesion initiation and progression in coronary arteries (*4*, *51*), adverse cardiovascular outcomes such as myocardial infarction (*52*, *53*), long-term restenosis after carotid bifurcation endarterectomy (*54*), aortic vascular stiffening (*55*), and coronary lesion phenotypes (*56*).

Several limitations should nevertheless be considered. Personalized CFD models inherently involve uncertainties and modeling assumptions—Newtonian blood rheology, rigid walls, idealized boundary conditions, etc. (*57*)—that may influence the topological analysis. However, the primary alternative approach, namely the direct *in vivo* measurement of velocity fields, vorticity, and WSS using 4D flow MRI, is likewise subject to known limitations and sources of inaccuracy (*58*, *59*). Despite these challenges, the 4D flow MRI–derived findings have remained fully consistent with the predictions of the theoretical topological framework, providing additional confidence in the robustness of the proposed methodology.

In conclusion, by translating principles from atmospheric physics to the cardiovascular domain, this study advances a topological perspective on cardiovascular flow science and introduces a unified approach for interpreting hemodynamic complexity. The proposed framework provides a mechanistic basis for understanding and quantifying “hemodynamic storms” in the brain, bridging physical fluid dynamics and vascular biomechanics within a common conceptual foundation. Future longitudinal studies tracking aneurysm morphology over time will be essential for determining whether this framework can help predict which aneurysms are most likely to grow and, ultimately, to rupture. More broadly, our findings suggest that both rotational (tornado-like) and nonrotational (downburst-like) fluid structures share a unified topological basis governing their formation and persistence under pathophysiological flow conditions, albeit with potentially distinct pathobiological implications. Although the framework is demonstrated here in cerebral aneurysms, selected as a paradigm of hemodynamic complexity, the general nature of the underlying topological principles suggests that it may be readily extended to other vascular districts and to a broader spectrum of cardiovascular diseases. By distilling complex hemodynamic data into interpretable spatiotemporal representations, this topological framework may facilitate a mechanistic understanding of cardiovascular disease progression and outcome, as supported by recent studies that have adopted WSS topology-based quantities as biomarkers (*4*, *51–54*). In this way, it lays the groundwork for studies aimed at establishing causal links between flow organization and cardiovascular diseases.

## MATERIALS AND METHODS

### Theoretical Remarks

Let Ω ⊂ ℝ^3^ be a bounded domain representing a fluid region enclosed by the material surface *δ*Ω — specifically, in our case, the lumen of a blood vessel. It is consolidated knowledge (*51*, *60*) that for a Newtonian fluid the tangential (***u****_t_*) and the normal (***u****_n_*) components of the fluid velocity in the near-wall region—expressed in a local coordinate system (**ζ_1_**, **ζ_2_**, ***n***) where **ζ_1_** and **ζ_2_**, are the orthogonal unit tangent vectors and ***n*** the unit normal vector to the wall—can be approximated in terms of the WSS, ***τ***(**ζ_1_**, **ζ_2_**), on the surface boundary *δ*Ω, as follows:

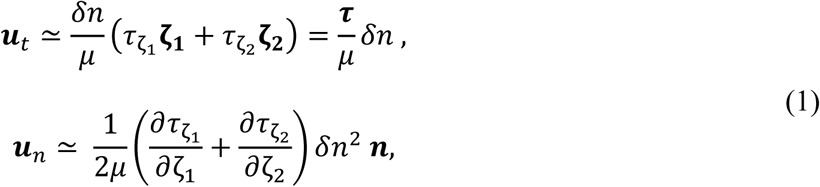

where *μ* is the dynamic viscosity of the fluid and *δn* is the intravascular normal distance from the wall (as already reported elsewhere (*60*), if the flow is unsteady, additional convective transport components of ***u****_t_* and ***u****_n_* in eq. (1) are an order of magnitude smaller and can be considered to have a lesser degree of importance in the current study). Since the normal component of the near-wall velocity is second order in *δn*, eq. (1) indicates that WSS on the surface serves as a direct marker of the velocity dynamics in the immediate vicinity of the vessel wall. The near-wall fluid dynamics can be further understood through the theoretical relationship linking the WSS, ***τ***, to the vorticity vector field, ***ω***, which is solenoidal in Ω (i.e., the condition ∇ ⋅ ***ω*** = 0, is always satisfied in Ω). In particular, the WSS is directly related to the SV ***ω*_δ_**_Ω_ (i.e., the vorticity defined on *δ*Ω, always tangent to the wall due to the no-slip condition; see Fig. 1B), through the equation (*9*):

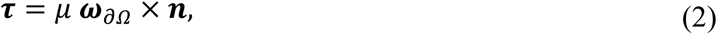

where ***n*** is the unit vector normal to *δ*Ω, oriented from the wall toward the fluid domain (for the derivation of eq. (2), see Supplementary Materials, Methods).

Very recently, it has been shown that the near-wall fluid dynamics can be further elucidated by analyzing a key fluid-mechanics quantity, the vorticity diffusion flux normal to the wall, denoted as *σ_n_* and defined as follows (*9*):

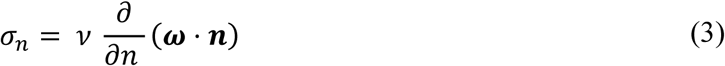

where *ν* is the kinematic viscosity of the fluid (for the derivation of eq. (3), see Supplementary Materials, Methods). The vorticity diffusion flux *σ_n_*—whose expression is derived by integrating the viscous term from the vorticity transport equation (see Supplementary Materials, Methods)— quantifies the amount of vorticity diffused into or out of the boundary *δ*Ω per unit area and unit time and provides a description of the kinematic evolution of the vorticity in the near-wall region (*9*, *61*). Eqs. (1) to (3) clearly reflect that the WSS topological skeleton acts as unambiguous indicator of both near-wall velocity dynamics and vorticity kinematics. This becomes particularly evident when the vorticity flux *σ_n_*, defined in eq. (3), is expressed in terms of curl of the WSS:

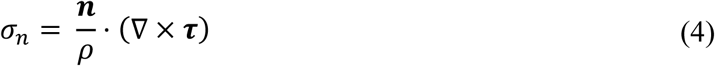

where *ρ* denotes the fluid density. The validity of eq. (4) has been demonstrated, for example, by Mazzi et al. (*9*), and its physical interpretation is that the tendency of WSS to rotate on *δ*Ω always implies a deflection of vortex lines from parallel to normal to the wall (or vice versa, depending on the direction of rotation for ***τ***) in the near-wall region.

Consequently, a detailed examination of these relationships provides valuable insights into the complex interactions that take place at the fluid-solid interface (in this case, at the blood vessel-wall interface). In particular, and as previously noted, a closer examination (which can be found in the Supplementary Materials, Methods) of eqs. (3) and (4) reveals two distinct scenarios by which existing near-wall vorticity interacts with the wall (*9*):

i. when *σ_n_* = 0. In this case, the gradient of ***ω*** in the direction normal to the wall vanishes. The resulting absence of vorticity diffusion flux across the wall implies that vortex lines near the wall remain aligned parallel to it, a configuration that can be represented topologically as shown in Fig. 1B;
ii. when *σ_n_* ≠ 0. The presence of a net vorticity diffusion flux in the normal direction causes vortex lines to deviate from parallel alignment, tilting either toward or away from the wall. This deflection of the near-wall vorticity field lines is invariably accompanied by a rotation of the WSS on *δ*Ω -a behavior consistent with eq. (2)-as illustrated in Fig. 1B.

Building on the relationship in eq. (2), which shows that a fixed point of ***τ*** is necessarily a fixed point of ***ω*_δ_**_Ω_ and vice versa, we recently proposed a theoretical taxonomy classifying the mathematically possible combinations of reciprocal stability properties and nearby field configurations for WSS and SV fixed points (*9*). The way this taxonomy was derived is presented in the Supplementary Materials, Methods (where a summary of the admissible relationships between fixed points of ***τ*** and ***ω*_δ_**_Ω_ in presence of zero or nonzero wall-normal vorticity diffusion flux, is provided in Supplementary Materials, Fig. S1).

Briefly, we demonstrated that, when *σ_n_* = 0 (*9*): (i) a node or star node for ***τ***, whether stable or unstable, always corresponds to a center for ***ω*_δ_**_Ω_—these fixed points are associated with non-rotational fluid structures emanating from the wall or impinging on the wall; (ii) a saddle point for ***τ*** always corresponds to a saddle point for ***ω*_δ_**_Ω_, and vice versa—these fixed points present both velocity and vortex lines parallel to the material surface, in the near wall (see Supplementary Materials, Methods). Contrarily, when *σ_n_* ≠ 0 (*9*): (i) a node or degenerate node for ***τ***, whether stable or unstable, can correspond to a (stable or unstable) node or degenerate node, or to a (stable or unstable) focus for ***ω*_δ_**_Ω_. In these cases, fixed points are associated with non-rotational fluid structures emanating from/impinging on the wall; (ii) a stable or unstable focus for ***τ*** can correspond to a (stable or unstable) node or degenerate node, or to a (stable or unstable) focus for ***ω*_δ_**_Ω_. In these cases, fixed points are associated with rotational fluid structures emanating from or impinging on the wall; (iii) a saddle point for ***τ*** always corresponds to a saddle point for ***ω*_δ_**_Ω_, and vice versa. Saddle points are unavoidably associated with non-rotational fluid structures, with velocity streamlines and vortex lines which are oppositely emanating from and impinging on the wall (see Supplementary Materials, Methods).

### Computational Hemodynamics

High-fidelity CFD simulations—where “high-fidelity” refers to the use of numerical schemes and discretization strategies designed to avoid the artificial suppression of high-frequency flow instabilities known to occur in aneurysms (*62*)—of three unruptured aneurysm cases from Toronto Western Hospital, representing stable (S), moderately unstable (Um) and highly unstable (Uh) saccular hemodynamics, were used to identify tornadic phenomena. Case Um (sac height 7.8 mm; sac width 7.4 mm) was an internal carotid artery sidewall aneurysm imaged by routine diagnostic 3D rotational angiography, for which a cycle-averaged inflow rate proportional to inlet area was assumed (*63*). Cases S (sac height 4.8 mm; sac width 4.7 mm) and Uh (sac height 7.9 mm; sac width 9.2 mm) were middle cerebral artery bifurcation aneurysms from a research study on 4D computed tomography, which allowed for individual cycle-averaged inflow rates to be measured (*64*). In all cases, pulsatile flow assumed the same population-averaged inflow waveform shape (*65*) shown in Fig. 3A, which was scaled by each case’s cycle-averaged inflow rate. Three-dimensional lumen segmentation was performed with a morphological gradient-based watershed algorithm2. Segmented lumen geometries and flow rates were de-identified and then shared for CFD purposes under University Health Network Research Ethics Board approvals 09-0059 (Case Um) and 16-53960BE (Cases S and Uh).

The procedures for geometry reconstruction and the meshing strategy, as well as CFD settings and their verification, have been described elsewhere (*8*, *66*, *67*). Briefly, tetrahedral meshes comprising 2.2-3.3 million elements, with a uniform density of 0.13 mm in the aneurysm sac and progressively fine layers towards the wall throughout, were generated using the Vascular Modelling Toolkit software. Further details on mesh verification and meshing heuristics are provided elsewhere (*67*, *68*). The Navier-Stokes equations were numerically solved using the finite element solver OASIS— a minimally dissipative and energy-preserving solver employing a fractional-step algorithm to decouple velocity and pressure (*69*). Simulations were second-order accurate in terms of spatial and temporal discretization. Blood was assumed as a homogeneous Newtonian fluid (*70*). Under the rigid wall assumption, pulsatile blood flow was simulated, with a typical temporal resolution of 10,000 time-steps per cardiac cycle, conservatively chosen to ensure satisfaction of well-known Courant-Friedrich-Lewy stability conditions for the fine meshes used. Models included sufficiently long segments of the tortuous parent artery upstream so that the necessary imposition of fully developed Womersley velocity profiles would be mitigated at the aneurysm sac. At the outlets, flow rates were distributed dynamically according to a previously validated flow-splitting technique (*71*).

Tornadic flow phenomena within aneurysmal sacs were visualized using instantaneous velocity streamlines and vortex lines. These features were derived from CFD data through forward and backward integration initiated in the vicinity of the fixed point on the luminal surface, enabling visualization of the associated local flow dynamics. Intra-aneurysmal vortical structures were further examined using swirling strength, a quantitative measure of local rotational motion within the flow. Swirling strength was defined as |*λ_ci_*|, representing the magnitude of the imaginary component of the complex conjugate eigenvalue of the velocity gradient tensor (*72*). To investigate vortices generated by the interaction of downburst-like fluid structures with the vessel wall, vortex corelines were also identified. Technically, a point is classified as belonging to a vortex coreline when the local velocity gradient tensor exhibits complex-conjugate eigenvalues and the local velocity vector is aligned with the real eigenvector of this tensor. Here, vortex corelines were identified using the canonical Parallel Vector Method (*73*, *74*).

From CFD data we also derived the two WSS-based quantities conventionally adopted to identify near-wall flow disturbances in cerebral aneurysms, namely the time-average WSS (TAWSS)— which is the average magnitude of WSS over the cardiac cycle at each point on the luminal surface—and the Oscillatory Shear Index (OSI), which is a measure of WSS flow reversal across the cardiac cycle (see Supplementary Materials, Methods).

### Topological Skeleton Analysis

At each time-step of the cardiac cycle, fixed points of WSS and SV on the luminal surface of CFD cerebral aneurysm models were identified by computing the Poincaré index (*75*), a topological invariant that takes the value -1 in the presence of a saddle point, +1 in the presence of a node, focus, or center, and 0 in regions free of fixed points. The Poincaré index enables localization of fixed points but does not provide complete information on their type or stability. These properties were therefore determined by computing and analyzing the eigenvalues of the Jacobian matrices of the WSS and SV fields, following an established methodology described in previous studies (*8*, *76*). Exhaustive details on the fixed-point classification procedure are provided in the Supplementary Materials (Methods, see table S1 for a summary).

Building on the Eulerian framework proposed in (*8*), the divergence of the normalized WSS field, DIV_WSS_, defined as:

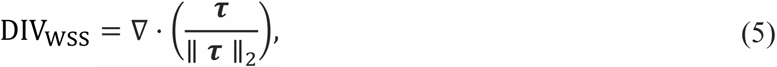

was used here to detect WSS manifolds on the surface of the aneurysmal sac (Fig. 1E). Application of volume contraction theory to the near-wall velocity field in eq. (1), showed that negative DIV_WSS_values approximate unstable WSS manifolds, whereas positive values approximate stable WSS manifolds (*8*, *51*).

In the present study, we adopted a fixed-point carousel visualization, a recently proposed strategy to visualize the four-dimensional nature of WSS fixed-point kinematics along the cardiac cycle (*23*). Briefly, this recently introduced topographic mesh parameterization strategy enables the reduction of the three-dimensional aneurysm sac anatomy to a two-dimensional plane, eliminating occlusion. Technically, the neck of the sac was delineated semi-automatically, and a least-squares conformal mapping (*77*) was applied to warp the sac onto a planar surface, analogous to cartographic mapping. Least-squares conformal mapping was specifically chosen to minimize length/area distortions, to preserve the Euclidean distances between instantaneous fixed-point locations and the relative surface areas of the sac dome, middle and neck regions. The resulting instantaneous two-dimensional maps with fixed points were then swept, in time, around a circle to reinforce the periodic nature of the cardiac cycle. Finally, a cut-away toroidal surface was added to provide spatiotemporal cues in the absence of otherwise-cluttering planar surfaces (Fig. 1E). Further rationales and methodological details for what we have termed the “fixed-point carousel” visualization are provided elsewhere (*23*). By leveraging the theoretical relationship between WSS and SV (*9*), the WSS fixed-point carousel enables precise temporal and spatial localization of tornadic fluid phenomena within the aneurysmal sac. This provides an intuitive and physically grounded means to characterize the onset, evolution, and lifetime of near-wall fluid structures, and was used to localize representative examples of the various tornadic phenomena.

Moreover, to quantify the residence time of instantaneous WSS fixed points at specific locations of the aneurysmal sac over the cardiac cycle, we adopted a recently proposed formulation that measures the fraction of the cycle during which a WSS fixed point resides within a generic mesh surface element. The instantaneous contributions to this residence time measure are weighted by the absolute value of the instantaneous WSS divergence, which reflects the local strength of the contraction or expansion action exerted by the shear forces (*8*):

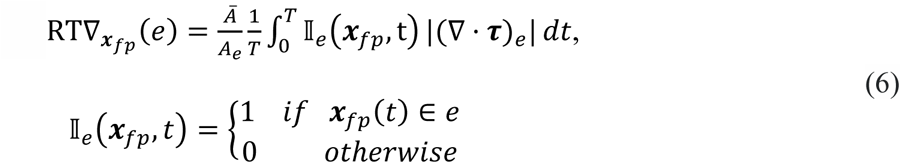

Here, ***x****_fp_*(*t*) denotes the position of the WSS fixed point at time *t* ∈ [0, *T*]; *e* is the generic triangular element of the luminal surface mesh with area *A_e_*; *Ā* is the average surface area of all triangles; and (∇ · ***τ***)*_e_* is the instantaneous WSS divergence evaluated in the surface triangle *e* containing ***x****_fp_*(*t*). As previously shown (*8*), eq. (6) accentuates the contribution of fixed points surrounded by strong local WSS contraction or expansion actions, even if these occur during a small portion of the cardiac cycle.

### 4D flow MRI Acquisition Protocol and Data Processing

Two patients with intracranial aneurysm were selected from a cohort of patients undergoing surveillance monitoring for known cerebral aneurysms at the University of California San Francisco hospital (UCSF). Patients with untreated cerebral aneurysms are monitored annually using non-invasive MRI. The two cases analyzed here include a cavernous internal carotid artery aneurysm (Case M1, sac height 4.4 mm and sac width 6.7 mm) and a middle cerebral artery bifurcation aneurysm (Case M2, sac height 5.6 mm and sac width 8.1 mm). MR Imaging was acquired with a 3T scanner (Siemens Magnetom Vida) using a 20-channel head and neck coil. Following administration of a gadolinium-based contrast agent (0.2 ml/kg, Dotarem), 4D Flow MRI was acquired in an axial-oblique 3D volume centered on the cerebral arteries of interest. Images were obtained using retrospective cardiac gating and compressed sensing with the following pulse sequence parameters: TR = 4.2 ms, TE = 2.8 ms, native spatial resolution = 1.0 x 1.0 x 1.0 mm^3^, reconstructed resolution = 0.5 x 0.5 x 1.0 mm^3^, flip angle = 12°, 22 cardiac phases, and velocity-encoding value of 100 cm/s.

4D flow data were post-processed using an in-house dedicated Python-based toolbox (*78*). Briefly, the 4D flow MRI data were corrected for background phase offsets, eddy currents, and velocity aliasing when needed. Magnitude and phase images were then combined to generate a phase-contrast angiogram (PCA). The main cerebral arteries forming the circle of Willis were segmented by applying a global threshold to the PCA. Threshold selection was guided by verification of flow conservation between parent arteries and downstream branch vessels. Time-resolved 3D vector phase velocity data and lumen boundary masks were de-identified and then shared for the purposes of extracting vortical structures and WSS vector fields, under UCSF IRB approval 10-03060. In accordance with recent consensus recommendations for 4D flow MRI (*79*), the three acquired phase-velocity data were interpolated onto a denser unstructured grid using a Gaussian kernel. Velocity gradients were subsequently computed on the interpolated field using cell shape functions evaluated at parametric coordinates. Flow features were visualized using multiple tools implemented in the open-source platform ParaView.

## Funding

D.G. and K.C. acknowledge the support of «ASSOCIATE» project (code 2022L7KK7L, CUP E53D23003480006.) – funded by European Union – Next Generation EU within the PRIN 2022 program (D.D. 104 -02/02/2022 Ministero dell’Università e della Ricerca). D.A.S acknowledges the support of a Discovery grant (RGPIN-2018-04649) from the Natural Sciences & Engineering Research Council of Canada. Priority core-hours for high-fidelity CFD were provided by SciNet, which is funded by: the Canada Foundation for Innovation; the Government of Ontario; Ontario Research Fund - Research Excellence; and the University of Toronto. U.M. acknowledges the support of «COMPUTES» project (code 2022ZKEP8SAC, CUP E53C24002930006), Next Generation EU within the PRIN 2022 program (D.D. 104 - 02/02/2022 Ministero dell’Università e della Ricerca).

## Competing interests

We have no conflicts of interest to disclose.

## Data, code, and materials availability

Data and analysis codes supporting the findings of the study have been deposited in the repository Figshare (*80*).

## Footnotes

1 Throughout, “tornadic” is used strictly in a topological and organizational sense rather than to imply identical physics.

2 Note that cases Um, S, and Uh correspond to cases A, B, and C from Natarajan et al. (23), respectively.

## Notes

https://doi.org/10.6084/m9.figshare.32270130

### Competing Interest Statement

The authors have declared no competing interest.

