## Supplementary Materials for "Evidence of tornadic phenomena in cerebral aneurysms"

*[1] PoliTo[BIO]Med Lab, Department of Mechanical and Aerospace Engineering, Politecnico di Torino, Turin, Italy; [2] Department of Biomedical Engineering, Georgia Institute of Technology, Atlanta, GA; [3] Department of Radiology and Biomedical Imaging, University of California San Francisco, San Francisco, California, USA; [4] Biomedical Simulation Laboratory, Department of Mechanical & Industrial Engineering, University of Toronto, Toronto, ON Canada*

### METHODS

#### Theoretical Remarks: Derivation of the Relationship between Wall Shear Stress and Surface Vorticity

In a domain $\Omega \subset \mathbb{R}^3$ where a Newtonian fluid flows, the wall shear stress (WSS) $\boldsymbol{\tau}$ on the surface boundary $\partial\Omega$ (i.e., the luminal surface of a vessel, Fig. 1B) can be defined as:

$$\boldsymbol{\tau} = (I - \boldsymbol{n} \otimes \boldsymbol{n}) \cdot (-2\,\mu\,\boldsymbol{n} \cdot \boldsymbol{S}_{\partial\Omega}), \qquad \text{(s1)}$$

where $I$ is the identity matrix, $\mu$ is the dynamic viscosity, $\boldsymbol{n}$ is the unit vector normal to $\partial\Omega$ (directed from the wall to the fluid), and $\boldsymbol{S}_{\partial\Omega}$ is the strain rate tensor on $\partial\Omega$, defined as:

$$\boldsymbol{S}_{\partial\Omega} = \frac{1}{2}(\nabla\boldsymbol{u}_{\partial\Omega} + \nabla\boldsymbol{u}_{\partial\Omega}^T). \qquad \text{(s2)}$$

In eqs. (s1) and (s2) the subscript $\partial\Omega$ denotes the restriction of a quantity on the luminal surface, and $\nabla\boldsymbol{u}_{\partial\Omega}$ is the velocity gradient tensor, that can be decomposed into the sum of its symmetric ($\boldsymbol{S}_{\partial\Omega}$) and skew-symmetric ($\boldsymbol{W}_{\partial\Omega}$) parts:

$$\nabla\boldsymbol{u}_{\partial\Omega} = \boldsymbol{S}_{\partial\Omega} + \boldsymbol{W}_{\partial\Omega} = \frac{1}{2}(\nabla\boldsymbol{u}_{\partial\Omega} + \nabla{\boldsymbol{u}_{\partial\Omega}}^T) + \frac{1}{2}(\nabla\boldsymbol{u}_{\partial\Omega} - \nabla{\boldsymbol{u}_{\partial\Omega}}^T). \qquad \text{(s3)}$$

It follows from eq. (s3) that tensor $\boldsymbol{S}_{\partial\Omega}$ can then be expressed as follows:

$$\boldsymbol{S}_{\partial\Omega} = \nabla\boldsymbol{u}_{\partial\Omega}^T + \boldsymbol{W}_{\partial\Omega}. \qquad \text{(s4)}$$

By substituting eq. (s4) in eq. (s1), the expression for $\boldsymbol{\tau}$ can be reformulated in the following terms:

$$\begin{aligned}\boldsymbol{\tau} &= (I - \boldsymbol{n} \otimes \boldsymbol{n}) \cdot \left[-2\,\mu\,\boldsymbol{n} \cdot \left(\nabla\boldsymbol{u}_{\partial\Omega}^T + \boldsymbol{W}_{\partial\Omega}\right)\right] \\ &= (I - \boldsymbol{n} \otimes \boldsymbol{n}) \cdot \left[2\,\mu\left(-\boldsymbol{n} \cdot \nabla\boldsymbol{u}_{\partial\Omega}^T - \boldsymbol{n} \cdot \boldsymbol{W}_{\partial\Omega}\right)\right].\end{aligned} \qquad \text{(s5)}$$

Because $\partial\Omega$ can be assumed to be rigid, the surface deformation stress, which can be expressed as $\boldsymbol{t}_s = -2\mu\boldsymbol{n}\cdot\left[(\nabla\cdot\boldsymbol{u}_{\partial\Omega})\mathbf{I} - \nabla\boldsymbol{u}_{\partial\Omega}^T\right]$, where $\mathbf{I}$ is the unit tensor, is zero (1). For an incompressible fluid such as blood, the condition $\nabla\cdot\boldsymbol{u}_{\partial\Omega} = 0$ holds, and therefore the term involving $(\nabla\cdot\boldsymbol{u}_{\partial\Omega})\mathbf{I}$ vanishes. It follows that the term $2\,\mu\,\boldsymbol{n}\cdot\nabla\boldsymbol{u}_{\partial\Omega}^T$ in eq. (s5) is also zero. Moreover, by exploiting the skew-symmetry of the spin tensor, eq. (s5) can be reformulated as:

$$\boldsymbol{\tau} = (I - \boldsymbol{n}\otimes\boldsymbol{n})\cdot[2\,\mu\,(-\boldsymbol{n}\cdot\boldsymbol{W}_{\partial\Omega})] = (I - \boldsymbol{n}\otimes\boldsymbol{n})\cdot[2\,\mu\,(\boldsymbol{W}_{\partial\Omega}\cdot\boldsymbol{n})]. \quad \text{(s6)}$$

For any generic vector $\boldsymbol{\xi}$, the spin tensor $\boldsymbol{W}_{\partial\Omega}$ satisfies the well-known relation (2):

$$2\boldsymbol{W}_{\partial\Omega}\cdot\boldsymbol{\xi} = \boldsymbol{\omega}_{\partial\Omega}\times\boldsymbol{\xi}, \quad \text{(s7)}$$

where $\boldsymbol{\omega}_{\partial\Omega}$ is the surface vorticity (SV), i.e., the vorticity on the luminal surface of the vessel. It follows that eq. (s6) can be reformulated as:

$$\begin{aligned}\boldsymbol{\tau} &= (I - \boldsymbol{n}\otimes\boldsymbol{n})\cdot(\mu\,\boldsymbol{\omega}_{\partial\Omega}\times\boldsymbol{n})\\ &= (\mu\,\boldsymbol{\omega}_{\partial\Omega}\times\boldsymbol{n}) - (\boldsymbol{n}\otimes\boldsymbol{n})\cdot(\mu\,\boldsymbol{\omega}_{\partial\Omega}\times\boldsymbol{n})\\ &= (\mu\,\boldsymbol{\omega}_{\partial\Omega}\times\boldsymbol{n}) - [(\mu\,\boldsymbol{\omega}_{\partial\Omega}\times\boldsymbol{n})\cdot\boldsymbol{n}]\boldsymbol{n}\\ &= \mu\,\boldsymbol{\omega}_{\partial\Omega}\times\boldsymbol{n},\end{aligned} \quad \text{(s8)}$$

where the term $(\mu\,\boldsymbol{\omega}_{\partial\Omega}\times\boldsymbol{n})\cdot\boldsymbol{n} = 0$, being vectors $(\mu\,\boldsymbol{\omega}_{\partial\Omega}\times\boldsymbol{n})$ and $\boldsymbol{n}$ orthogonal by construction.

A consequence of eq. (s8) is that the pair $(\boldsymbol{\tau}, \boldsymbol{\omega}_{\partial\Omega})$ is orthogonal on $\partial\Omega$. Moreover, based on eq. (s8) the WSS can be interpreted not only in terms of near-wall velocity dynamics (3), but also in terms of vorticity, dictated by the shear viscosity and the no-slip condition (2).

**Theoretical Remarks: Derivation of the Vorticity Diffusion Flux Normal to a Material Surface**

Technically, the expression adopted here for the vorticity diffusion flux normal to a material surface follows the formulation introduced by Lighthill for solid boundaries (4). Lighthill defined the boundary vorticity flux by analogy with the heat flux in Fourier's law. This definition can be justified by integrating the viscous term of the vorticity transport equation, $\nu\,\nabla^2\boldsymbol{\omega}$, where $\nu$ is the kinematic viscosity, over a control volume $V$ bounded by $\partial V$:

$$\int_V \nu\,\nabla^2\boldsymbol{\omega}\,dV = \oint_{\partial V} \nu\,\boldsymbol{n}\cdot\nabla\boldsymbol{\omega}\,dS \quad \text{(s9)}$$

In the present case, the vorticity diffusion flux normal to $\partial\Omega$—arising from the viscous term in the vorticity transport equation (5) —quantifies the amount of vorticity diffusing in or out of $\partial\Omega$ per unit area and unit time and, based on eq. (s9), can be expressed as:

$$\sigma_n = \nu\,\frac{\partial}{\partial n}(\boldsymbol{\omega}\cdot\boldsymbol{n}). \quad \text{(s10)}$$

For an incompressible fluid, vorticity is solenoidal, so that $\nabla\cdot\boldsymbol{\omega} = 0$ throughout $\Omega$. Because the velocity field satisfies the no-slip condition, $\boldsymbol{\omega}_{\partial\Omega}$ is tangent to $\partial\Omega$. Introducing a local coordinate system $(\boldsymbol{\zeta}_1, \boldsymbol{\zeta}_2, \boldsymbol{n})$, with $\boldsymbol{\zeta}_1$ and $\boldsymbol{\zeta}_2$ tangent to $\partial\Omega$ and $\boldsymbol{n}$ always normal to it, the solenoidal condition becomes:

$$\frac{\partial}{\partial n}(\boldsymbol{\omega}\cdot\boldsymbol{n})+\frac{\partial}{\partial \zeta_1}(\boldsymbol{\omega}\cdot\boldsymbol{\zeta_1})+\frac{\partial}{\partial \zeta_2}(\boldsymbol{\omega}\cdot\boldsymbol{\zeta_2})=0. \qquad \text{(s11)}$$

Since $\boldsymbol{\omega}_{\partial\Omega}$ lies on $\partial\Omega$, the vorticity diffusion flux normal to $\partial\Omega$ in eq. (s10) may therefore be reformulated in terms of surface derivatives as:

$$\sigma_n = \nu\,\frac{\partial}{\partial n}(\boldsymbol{\omega}\cdot\boldsymbol{n}) = -\nu\left[\frac{\partial}{\partial \zeta_1}(\boldsymbol{\omega}\cdot\boldsymbol{\zeta_1})+\frac{\partial}{\partial \zeta_2}(\boldsymbol{\omega}\cdot\boldsymbol{\zeta_2})\right] = -\nu\,\nabla_\pi\cdot\boldsymbol{\omega}, \qquad \text{(s12)}$$

where $\nu$ is the kinematic viscosity and $\nabla_\pi$ denotes the surface gradient operator.
In addition, a nonzero wall-normal vorticity diffusion flux $\sigma_n$ can be related to the curl of the WSS on $\partial\Omega$. Specifically, as already shown elsewhere (2), it can be easily demonstrated that:

$$\sigma_n = -\nu\,\nabla_\pi\cdot\boldsymbol{\omega} = \frac{\boldsymbol{n}}{\rho}\cdot(\nabla\times\boldsymbol{\tau}), \qquad \text{(s13)}$$

where $\rho$ is the fluid density.

### Topological Skeleton Analysis

After the identification of fixed points on the luminal surface of the aneurysm using a topological invariant, the Poincaré index (6), fixed-point type and stability classification was performed by analyzing the eigenvalues $\lambda_i$ of the Jacobian matrix $J$ of the WSS and SV fields. Because the orthogonal pair $(\boldsymbol{\tau}, \boldsymbol{\omega}_{\partial\Omega})$ lies on $\partial\Omega$, the fixed-point analysis was restricted to two dimensions, as reported in Table S1.

**Table S1**. Classification of fixed points for a two-dimensional field based on the eigenvalues $\lambda_i$ ($i$ = 1, 2) of the Jacobian matrix $J$. $I$ denotes the identity matrix.

| Eigenvalues | Fixed point |
|---|---|
| $\lambda_1 < 0 < \lambda_2$ | Saddle point |
| $\lambda_1, \lambda_2 > 0$ | Unstable node |
| $\lambda_1, \lambda_2 < 0$ | Stable node |
| $\lambda_1 = \lambda_2 > 0, J = \lambda I$ | Unstable star node |
| $\lambda_1 = \lambda_2 < 0, J = \lambda I$ | Stable star node |
| $\lambda_1 = \lambda_2 > 0, J \neq \lambda I$ | Unstable degenerate node |
| $\lambda_1 = \lambda_2 < 0, J \neq \lambda I$ | Stable degenerate node |
| $\lambda_{1,2} = \alpha \pm \beta i, \alpha, \beta \in \mathbb{R}^+$ | Unstable focus |
| $\lambda_{1,2} = -\alpha \pm \beta i, \alpha, \beta \in \mathbb{R}^+$ | Stable focus |
| $\lambda_{1,2} = \pm\beta i, \beta \in \mathbb{R}^+$ | Center |

In general, the eigenvalues $\lambda_\mathrm{i}$ of the Jacobian matrix $J$ on $\partial\Omega$ are the roots of the characteristic polynomial $p_J(\lambda) = \det(J - \lambda I)$, which can be expressed in terms of trace and determinant as:

$$\lambda^2 - \mathrm{tr}(J)\lambda + \det(J) = 0, \qquad \text{(s14)}$$

where $\mathrm{tr}(J) = \lambda_1 + \lambda_2$ and $\det(J) = \lambda_1\lambda_2$. As shown in Fig. 1D of the main text, fixed points can be classified based on $\mathrm{tr}(J)$ and $\det(J)$. Specifically, if $\det(J) < 0$, the fixed point is a saddle point, otherwise it can be a node, a focus or a center. Particularly, if $\det(J) > \frac{\mathrm{tr}^2(J)}{4}$, the fixed point is a focus. If $\det(J) < \frac{\mathrm{tr}^2(J)}{4}$ , the fixed point is a node. If $\det(J) = \frac{\mathrm{tr}^2(J)}{4}$, the fixed point is either star or a degenerate node. Finally, if $\mathrm{tr}(J) = 0$, the fixed point is a center. For node- and focus-type configurations, stability is determined by the sign of $\mathrm{tr}(J)$: $\mathrm{tr}(J) < 0$ identifies a stable fixed point, whereas $\mathrm{tr}(J) > 0$ identifies an unstable one.

According to eq. (s8), in the local coordinate system $(\boldsymbol{\zeta_1}, \boldsymbol{\zeta_2}, \boldsymbol{n})$, the WSS can be expressed in terms of the SV components $\boldsymbol{\omega}_{\partial\Omega} = \left(\omega_{\zeta_1}, \omega_{\zeta_2}, 0\right)$ as:

$$\boldsymbol{\tau} = \left(\tau_{\zeta_1}, \tau_{\zeta_2}, 0\right) = \mu \boldsymbol{\omega}_{\partial\Omega} \times \boldsymbol{n} = \mu \det \begin{pmatrix} \boldsymbol{\zeta_1} & \boldsymbol{\zeta_2} & \boldsymbol{n} \\ \omega_{\zeta_1} & \omega_{\zeta_2} & 0 \\ 0 & 0 & 1 \end{pmatrix} = (\mu\omega_{\zeta_2}, -\mu\omega_{\zeta_1}, 0). \qquad \text{(s15)}$$

It follows that:

$$\forall\, \tilde{\boldsymbol{x}} \in \partial\Omega : \ \boldsymbol{\tau}(\tilde{\boldsymbol{x}}) = 0 \Leftrightarrow \boldsymbol{\omega}_{\partial\Omega}(\tilde{\boldsymbol{x}}) = 0, \qquad \text{(s16)}$$

that is, a point $\tilde{\boldsymbol{x}} \in \partial\Omega$ is a fixed point of $\boldsymbol{\tau}$ if and only if it is also a fixed point of $\boldsymbol{\omega}_{\partial\Omega}$. However, although $\boldsymbol{\tau}$ and $\boldsymbol{\omega}_{\partial\Omega}$ share the same fixed-point locations, the type and stability of the corresponding fixed point may differ.

In the local coordinate system $(\boldsymbol{\zeta_1}, \boldsymbol{\zeta_2})$ tangent to $\partial\Omega$, consider the case in which, at point $\tilde{\boldsymbol{x}} \in \partial\Omega$,

$$\sigma_n = -\nu\, (\nabla_\pi \cdot \boldsymbol{\omega})_{\tilde{\boldsymbol{x}}} = 0. \qquad \text{(s17)}$$

It follows that, at any fixed point $\tilde{\boldsymbol{x}} \in \partial\Omega$ such that $\boldsymbol{\tau}(\tilde{\boldsymbol{x}}) = \boldsymbol{\omega}_{\partial\Omega}(\tilde{\boldsymbol{x}}) = 0$ and $\sigma_n = 0$, the following relation holds:

$$\frac{\partial\omega_{\zeta_1}}{\partial\zeta_1} = -\frac{\partial\omega_{\zeta_2}}{\partial\zeta_2}. \qquad \text{(s18)}$$

The characteristic polynomial of the Jacobian matrix $J$ of $\boldsymbol{\tau}$, evaluated at $\tilde{\boldsymbol{x}} \in \partial\Omega$ can be expressed as:

$$p_{J(\boldsymbol{\tau})_{\tilde{\boldsymbol{x}}}}(\lambda) = \lambda^2 - \mathrm{tr}(J(\boldsymbol{\tau})_{\tilde{\boldsymbol{x}}})\lambda + \det(J(\boldsymbol{\tau})_{\tilde{\boldsymbol{x}}}) = 0, \qquad \text{(s19)}$$

where

$$J(\boldsymbol{\tau})_{\tilde{\boldsymbol{x}}} = \begin{pmatrix} \frac{\partial\tau_{\zeta_1}}{\partial\zeta_1} & \frac{\partial\tau_{\zeta_1}}{\partial\zeta_2} \\ \frac{\partial\tau_{\zeta_2}}{\partial\zeta_1} & \frac{\partial\tau_{\zeta_2}}{\partial\zeta_2} \end{pmatrix}_{\tilde{\boldsymbol{x}}} = \mu \begin{pmatrix} \frac{\partial\omega_{\zeta_2}}{\partial\zeta_1} & \frac{\partial\omega_{\zeta_2}}{\partial\zeta_2} \\ -\frac{\partial\omega_{\zeta_1}}{\partial\zeta_1} & -\frac{\partial\omega_{\zeta_1}}{\partial\zeta_2} \end{pmatrix}_{\tilde{\boldsymbol{x}}}, \qquad \text{(s20)}$$

By virtue of eq. (s18), $J(\boldsymbol{\tau})_{\tilde{\boldsymbol{x}}}$ is symmetric.

Similarly, because $\mathrm{tr}(J(\boldsymbol{\omega}_{\partial\Omega})_{\tilde{\boldsymbol{x}}}) = (\nabla \cdot \boldsymbol{\omega}_{\partial\Omega}) = (\nabla_\pi \cdot \boldsymbol{\omega}) = 0$, the characteristic polynomial of $J(\boldsymbol{\omega}_{\partial\Omega})$ evaluated at the same fixed point $\tilde{\boldsymbol{x}}$, becomes

$$p_{J(\boldsymbol{\omega}_{\partial\Omega})_{\tilde{\boldsymbol{x}}}} = \lambda^2 + \det(J(\boldsymbol{\omega}_{\partial\Omega})_{\tilde{\boldsymbol{x}}}) = 0. \tag{s21}$$

Moreover, since $\det(J(\boldsymbol{\omega}_{\partial\Omega})) = \frac{\det(J(\boldsymbol{\tau}))}{\mu^2}$, and $\det(J) = \lambda_1\lambda_2$, the following cases arise:

- If $\det(J(\boldsymbol{\omega}_{\partial\Omega})_{\tilde{\boldsymbol{x}}}) = \frac{\det(J(\boldsymbol{\tau})_{\tilde{\boldsymbol{x}}})}{\mu^2} < 0$, then the eigenvalues $\lambda_1$ and $\lambda_2$ are real and of opposite sign. Therefore, the fixed point $\tilde{\boldsymbol{x}}$ is a saddle point for both $\boldsymbol{\omega}_{\partial\Omega}$ and $\boldsymbol{\tau}$.
- If $\det(J(\boldsymbol{\omega}_{\partial\Omega})_{\tilde{\boldsymbol{x}}}) = \frac{\det(J(\boldsymbol{\tau})_{\tilde{\boldsymbol{x}}})}{\mu^2} > 0$, then the eigenvalues $\lambda_1$ and $\lambda_2$ of $J(\boldsymbol{\omega}_{\partial\Omega})_{\tilde{\boldsymbol{x}}}$ are purely imaginary, whereas those of $J(\boldsymbol{\tau})_{\tilde{\boldsymbol{x}}}$ are real and have the same sign. Therefore, the fixed point $\tilde{\boldsymbol{x}}$ is a center for $\boldsymbol{\omega}_{\partial\Omega}$ and a node (if $\det(J(\boldsymbol{\tau})_{\tilde{\boldsymbol{x}}}) < \frac{\mathrm{tr}^2(J(\boldsymbol{\tau})_{\tilde{\boldsymbol{x}}})}{4}$) or a star node (if $(\det(J(\boldsymbol{\tau})_{\tilde{\boldsymbol{x}}}) = \frac{\mathrm{tr}^2(J(\boldsymbol{\tau})_{\tilde{\boldsymbol{x}}})}{4}$) for $\boldsymbol{\tau}$.

In the same local coordinate system $(\boldsymbol{\zeta_1}, \boldsymbol{\zeta_2})$ , consider the case in which, at a point $\hat{\boldsymbol{x}} \in \partial\Omega$,

$$\sigma_n = -\nu\,(\nabla_\pi \cdot \boldsymbol{\omega})_{\hat{\boldsymbol{x}}} \neq 0. \tag{s22}$$

It then follows that, at any fixed point $\hat{\boldsymbol{x}} \in \partial\Omega$ such that $\boldsymbol{\tau}(\hat{\boldsymbol{x}}) = \boldsymbol{\omega}_{\partial\Omega}(\hat{\boldsymbol{x}}) = 0$ and $\sigma_n \neq 0$, the normal derivative of the vorticity component normal to $\partial\Omega$ in $\hat{\boldsymbol{x}}$ satisfies the following equation:

$$\frac{\partial}{\partial n}(\boldsymbol{\omega} \cdot \boldsymbol{n}) = -\frac{\partial \omega_{\zeta_1}}{\partial \zeta_1} - \frac{\partial \omega_{\zeta_2}}{\partial \zeta_2} \neq 0. \tag{s23}$$

The characteristic polynomials of the $J$ matrices of $\boldsymbol{\tau}$ and $\boldsymbol{\omega}_{\partial\Omega}$, evaluated at $\hat{\boldsymbol{x}}$, are:

$$p_{J(\boldsymbol{\tau})_{\hat{\boldsymbol{x}}}}(\lambda) = \lambda^2 - \mathrm{tr}(J(\boldsymbol{\tau})_{\hat{\boldsymbol{x}}})\lambda + \det(J(\boldsymbol{\tau})_{\hat{\boldsymbol{x}}}) = 0, \tag{s24}$$

and

$$p_{J(\boldsymbol{\omega}_{\partial\Omega})_{\hat{\boldsymbol{x}}}}(\lambda) = \lambda^2 - \mathrm{tr}(J(\boldsymbol{\omega}_{\partial\Omega})_{\hat{\boldsymbol{x}}})\lambda + \det(J(\boldsymbol{\omega}_{\partial\Omega})_{\hat{\boldsymbol{x}}}) = 0. \tag{s25}$$

In this case $J(\boldsymbol{\tau})_{\hat{\boldsymbol{x}}}$ is no longer a symmetric matrix, because $\frac{\partial \omega_{\zeta_1}}{\partial \zeta_1} + \frac{\partial \omega_{\zeta_2}}{\partial \zeta_2} = \nabla_\pi \cdot \boldsymbol{\omega} \neq 0$.

Moreover, since $\det(J(\boldsymbol{\omega}_{\partial\Omega})) = \frac{\det(J(\boldsymbol{\tau}))}{\mu^2}$, the following cases arise:

- if $\det(J(\boldsymbol{\omega}_{\partial\Omega})_{\hat{\boldsymbol{x}}}) = \frac{\det(J(\boldsymbol{\tau})_{\hat{\boldsymbol{x}}})}{\mu^2} < 0$, then eigenvalues $\lambda_1$ and $\lambda_2$ are real numbers but with different sign and thus $\hat{\boldsymbol{x}}$ is a saddle point for both $\boldsymbol{\omega}_{\partial\Omega}$ and $\boldsymbol{\tau}$;

- if $\det(J(\boldsymbol{\omega}_{\partial\Omega})_{\hat{\boldsymbol{x}}}) = \frac{\det(J(\boldsymbol{\tau})_{\hat{\boldsymbol{x}}})}{\mu^2} > 0$, then eigenvalues $\lambda_1$ and $\lambda_2$ may be either complex conjugates or real numbers with the same sign. Accordingly, $\hat{\boldsymbol{x}}$ may correspond to a focus, a node, or a degenerate node fixed point in each field, with the specific classification determined separately from the corresponding trace and determinant (as shown in Fig. 1D of the main text).

A summary of the admissible relationships between fixed points of WSS and SV in the presence of zero and nonzero vorticity diffusion flux normal to the wall is provided in Fig. S1.

**Figure S1** Theoretically admissible relationships between WSS and SV fixed points in the presence of zero and nonzero vorticity diffusion flux normal to the wall.

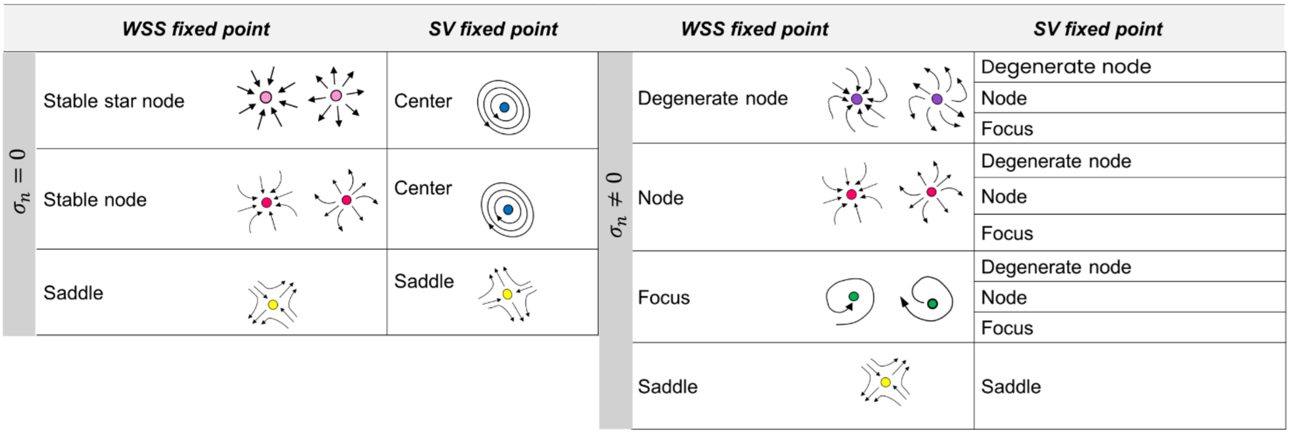


#### Conventional Wall Shear Stress-based Quantities

In addition to WSS topological features, the two WSS-based quantities conventionally adopted to identify near-wall flow disturbances in cerebral aneurysms, namely the time-averaged WSS (TAWSS) and the Oscillatory Shear Index (OSI) were computed:

$$\text{TAWSS} = \frac{1}{T}\int_0^T \|\, \boldsymbol{\tau} \,\|_2 \, dt$$

$$\text{OSI} = \frac{1}{2}\left(1 - \frac{\|\int_0^T \boldsymbol{\tau}\, dt\|_2}{\int_0^T \|\boldsymbol{\tau}\|_2 dt}\right) \quad 0 \leq \text{OSI} \leq 0.5 \tag{s26}$$

where $T$ is the cardiac cycle duration. In cerebral aneurysms, (i) low TAWSS regions are often associated with disturbed, slow fluid motion and have been linked to inflammatory wall remodeling and possible weakening, (ii) high TAWSS can relate to strong impingement jets and potential wall damage, and (iii) high OSI is often associated with unstable flow patterns, which are thought to contribute to aneurysm growth or rupture risk, typically in combination with low TAWSS.

**MOVIE 1.** Hemodynamics over the cardiac cycle of CASE S (characterized by mostly stable flow), visualized in terms of instantaneous velocity magnitude, isosurfaces of swirling strength ($\lambda_{ci} = 0.75\, s^{-1}$), and instantaneous WSS fixed points and manifolds on the surface of the aneurysmal sac. WSS unstable and stable manifolds are identified by negative (-ve) and positive (+ve) values of the divergence of normalized WSS ($DIV_{WSS}$), respectively.

**MOVIE 2.** Hemodynamics over the cardiac cycle of CASE Um (characterized by moderately unstable flow), visualized in terms of instantaneous velocity magnitude, isosurfaces of swirling strength ($\lambda_{ci} = 0.4\, s^{-1}$), and instantaneous WSS fixed points and manifolds on the surface of the aneurysmal sac. WSS unstable and stable manifolds are identified by negative (-ve) and positive (+ve) values of the divergence of normalized WSS ($DIV_{WSS}$), respectively.

**MOVIE 3.** Hemodynamics over the cardiac cycle of CASE Uh (characterized by highly unstable flow), visualized in terms of instantaneous velocity magnitude, isosurfaces of swirling strength ($\lambda_{ci} = 0.8\, s^{-1}$), and instantaneous WSS fixed points and manifolds on the surface of the aneurysmal sac. WSS unstable and stable manifolds are identified by negative (-ve) and positive (+ve) values of the divergence of normalized WSS ($DIV_{WSS}$), respectively.

**MOVIE 4.** *In vivo* 4D flow MRI visualization of blood flow in Cases M1 and M2. Instantaneous velocity streamlines, color-coded by velocity magnitude, are shown over the cardiac cycle.
